# Substrate access mechanism in a novel membrane-bound phospholipase A of *Pseudomonas aeruginosa* concordant with specificity and regioselectivity

**DOI:** 10.1101/2021.06.29.450291

**Authors:** Sabahuddin Ahmad, Christoph Heinrich Strunk, Stephan N. Schott-Verdugo, Karl-Erich Jaeger, Filip Kovacic, Holger Gohlke

## Abstract

PlaF is a cytoplasmic membrane-bound phospholipase A_1_ from *Pseudomonas aeruginosa* that alters the membrane glycerophospholipid (GPL) composition and fosters the virulence of this human pathogen. PlaF activity is regulated by a dimer-to-monomer transition followed by tilting of the monomer in the membrane. However, how substrates reach the active site and how the characteristics of the active site tunnels determine the activity, specificity, and regioselectivity of PlaF for natural GPL substrates has remained elusive. Here, we combined unbiased and biased all-atom molecular dynamics (MD) simulations and configurational free energy computations to identify access pathways of GPL substrates to the catalytic center of PlaF. Our results map out a distinct tunnel through which substrates access the catalytic center. PlaF variants with bulky tryptophan residues in this tunnel revealed decreased catalysis rates due to tunnel blockage. The MD simulations suggest that GPLs preferably enter the active site with the *sn*-1 acyl chain first, which agrees with the experimentally demonstrated PLA_1_ activity of PlaF. We propose that the acyl chain-length specificity of PlaF is determined by the structural features of the access tunnel, which results in favorable free energy of binding of medium-chain GPLs. The suggested egress route conveys fatty acid products to the dimerization interface and, thus, contributes to understanding the product feedback regulation of PlaF by fatty acid-triggered dimerization. These findings open up opportunities for developing potential PlaF inhibitors, which may act as antibiotics against *P. aeruginosa*.

## Introduction

*Pseudomonas aeruginosa* is an opportunistic and versatile pathogen, which causes infections in plants [1] and humans [2]. It is a multi-drug resistant Gram-negative bacterium and a frequent cause of nosocomial infections [3]. The pathogenicity of *P. aeruginosa* relies on both cell-associated and extracellular virulence factors [3]. Among those virulence factors are phospholipases [4, 5], including phospholipase A_1_ (PLA_1_), which hydrolyze cellular glycerophospholipids (GPLs) at the *sn*-1 position into lysoglycerophospholipids (LGPLs) and fatty acids (FAs) [6, 7].

GPLs primarily form bilayers, which maintain a permeability barrier for cells and organelles [8], while membrane-bound LGPLs can destabilize membrane integrity in Gram-negative bacteria [9, 10]. GPLs [11] and LGPLs [12, 13] can regulate the function and stability of membrane proteins. Interestingly, biofilm formation and growth phase transitions in *P. aeruginosa* are accompanied by the alteration of membrane GPL composition [14, 15]. FAs belong to the diffusible signal factor family (DSF) and are possible signal molecules because they can diffuse through cell membranes and contribute to the regulation of diverse biological functions in various Gram-negative pathogens [16]. In *P. aeruginosa,* DSFs promote biofilm formation and antibiotic resistance [17, 18].

We recently identified PlaF, an integral, inner membrane PLA_1_ that has a profound role for membrane GPL remodeling in *P. aeruginosa*. Furthermore, a *P. aeruginosa* Δ*plaF* knockout strain showed strongly attenuated virulence in *Galleria mellonella* and human macrophages models compared to the wild-type, which suggests that PlaF-mediated GPL remodeling contributes to the virulence of *P. aeruginosa* [19]. Crosslinking (*in vivo* and *in vitro*) and micro-scale thermophoresis experiments showed that PlaF exists in both monomeric and dimeric configurations, although it is active only in the monomeric state [19]. The crystal structure of PlaF revealed that a homodimer is formed by interactions between the transmembrane (TM) and juxtamembrane (JM) regions [19]. The homodimer contains co-crystallized endogenous ligands, myristic acid (MYR) and undecanoic acid (UND) from *P. aeruginosa*, (Figure 1), which are non-covalently bound in the active site cavity [19]. Moreover, a complex T-shaped active site pocket formed by the TM, JM, and catalytic domain revealed three openings, one at the dimer interface, one close to the catalytic serine (S137), and one most likely pointing towards the membrane [19]. Molecular simulations of PlaF activation revealed a mechanism that involves a dimer-to-monomer transition followed by tilting of the monomer in the membrane [19]. The tilting orients PlaF in a configuration relative to the membrane such that substrates can directly access the cleft (Figure 1). By contrast, in the configuration observed in the crystal structure, the residues lining the opening of the active site cleft are more than 5 Å above the membrane surface (Figure 1). However, it is unknown how substrates reach the active site and how the characteristics of the active site determine the activity, specificity, and regioselectivity of PlaF for medium-chain GPLs.

**Figure 1:**
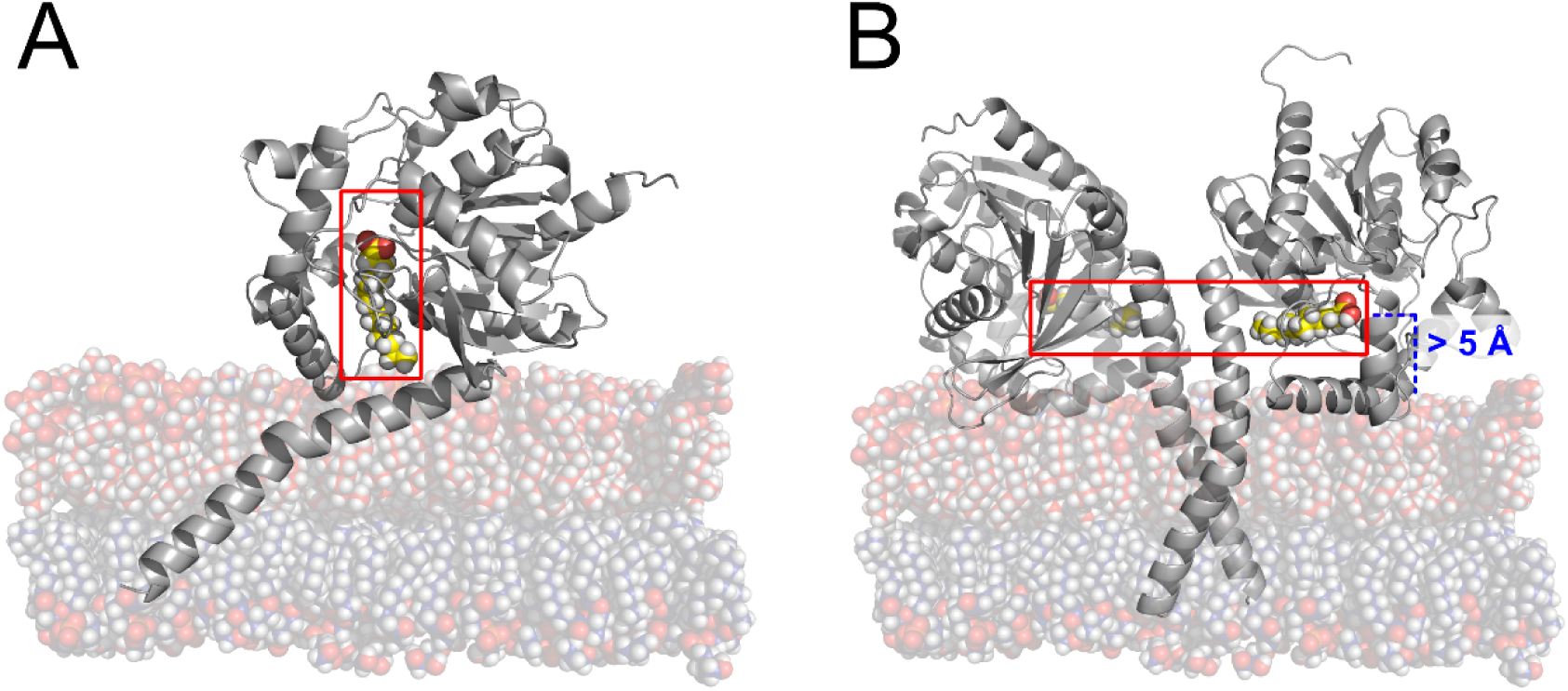
Schematic representation of the orientation of PlaF in the membrane. A) Chain A of dimeric PlaF in the tilted state; this state allows direct contact of the active site tunnel (red box) with the membrane. Yellow spheres represent the C atoms of the co-crystallized PlaF product, MYR. B) In dimeric PlaF, the active site tunnel is located > 5 Å above the membrane. Yellow spheres represent the C atoms of the co-crystallized PlaF products, MYR (left) and UND (right), within the active site tunnel.

The molecular mechanism underlying the access and binding of GPL to PLA is poorly understood in general because only a few PLA structures from microorganisms have been resolved, which either revealed closed conformations of their phospholipase domains [4, 20] or an accessible pocket that is predominantly hydrophobic [21] or amphipathic [22]. For the latter, regioselectivity was suggested to be achieved through binding of the GPL phosphate group to the polar pocket, which constrains the *sn*-1 acyl chain in a neighboring hydrophobic pocket [22]. Finally, structural analysis of the outer membrane PLA (OMPLA) from *Escherichia coli* in the complex with an inhibitor provided information about GPL recognition by this PLA [23, 24]. However, OMPLA is an integral β-barrel protein with a hydrophobic GPL binding cleft and the active site located at the β-barrel exterior. Hence, the mechanism by which PlaF recognizes GPL substrates must be conceptually different from that of OMPLA.

Here, we combined unbiased and biased all-atom molecular dynamics (MD) simulations and configurational free energy computations to identify access pathways to the catalytic center of PlaF. The results were validated by mutational and enzymatic studies on PlaF variants with blocked substrate access. Our results map out a distinct tunnel for substrate access within PlaF, provide explanations for the substrate specificity and PLA_1_ activity of PlaF, and suggest egress routes for hydrolysis products. These findings enhance our understanding of the mechanism by which membrane protein function is regulated through protein-GPL interactions.

## Results

### Access pathways to the catalytic site in PlaF

The crystal structure of PlaF revealed three pronounced tunnels, forming a large, T-shaped active site cleft. This cleft is compatible with binding bulky GPL substrates [19]. However, the structural dynamics of biomolecules may lead to variations in the tunnel shape [25]. Therefore, we reanalyzed trajectories from 10 replicas of unbiased MD simulations of 2 µs length for each of the systems di-PlaF (dimeric PlaF), PlaF_A_ (chain A from the crystal structure), PlaF_B_ (chain B from the crystal structure), and t-PlaF_A_ (chain A from the crystal structure in a tilted orientation) from our previous work [19] using CAVER [26]. CAVER analyzes and visualizes tunnels and channels in protein structures.

We primarily focus on t-PlaF_A_ because the tilted structure is likely the catalytically active form [19]. We identified the three tunnels that connect the active site of t-PlaF_A_ to its surface like in the crystal structure (Figure 2) [19]: Tunnel 1 (T1) and tunnel 2 (T2) point towards the membrane, and tunnel 3 (T3) opens to the periplasmic space > 15 Å above the membrane (Figure 2). T1 and T2 converge close to the active site and connect to T3. In the crystal structure, T1 contains MYR (chain A) and UND (chain B), which are hydrolysis products of GPL substrates with C14 and C11 acyl chain(s), respectively.

**Figure 2:**
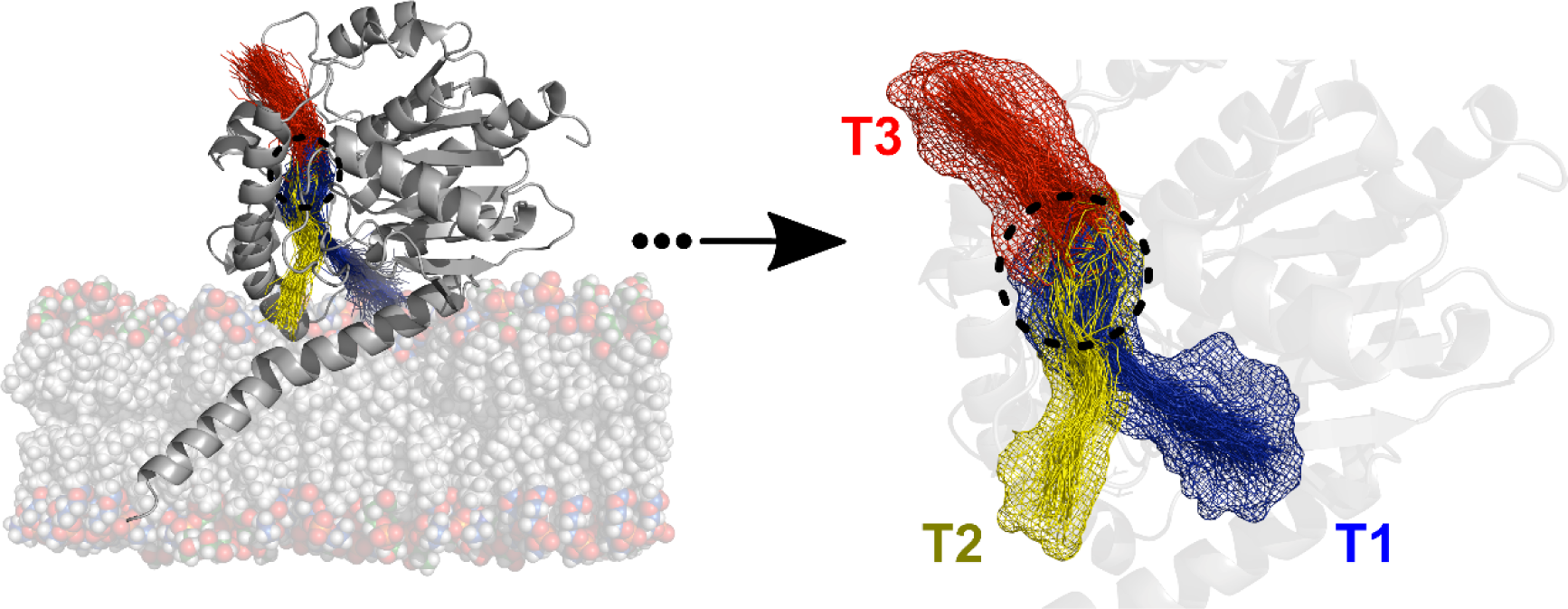
Clusters of tunnels identified in t-PlaF_A_ ensembles. Three major tunnel clusters connect the catalytic site (black dashed circle) of PlaF to the protein surface. Tunnels T1 and T2 point towards the membrane; tunnel T3 is located > 15 Å above the membrane, with its opening pointing into the periplasmic space.

T1 is the longest tunnel (Table 1) and was open more often than the other two tunnels (Table 1). The tunnel radii fluctuate between 2 Å and 5 Å depending on the location in the tunnel and the simulation length (Figure S1). The average bottleneck (narrowest part of the tunnel) radius of all tunnels is 2.26 ± 0.02 Å (mean ± standard error of the mean), which is close to the radius of glycerol (2.74 Å) [27], an essential component of all GPLs, but smaller than the radius of 1,2-dilauroyl-*sn*-glycero-3-phosphoglycerol (DLPG) (∼4.4 Å) deduced from the lipid’s area-per-lipid [28]. For comparison, tunnels in monomeric PlaF_A_, PlaF_B_, and the two chains of di-PlaF show open occurrences of ∼20% to ∼5% (Table S1), indicating no marked differences between monomeric and di-PlaF.

**Table 1:** Characteristics of tunnel clusters identified from unbiased MD simulations of t-PlaF_A_ using CAVER.

| Tunnel cluster | Occurrence <sup>a,b</sup> | Maximum bottleneck radius<sup>c</sup> | Average bottleneck radius<sup>c</sup> | Average length<sup>c</sup> |
| --- | --- | --- | --- | --- |
| T1 | 30.45 | 3.18 | 2.28 | 27.08 |
| T2 | 21.80 | 2.95 | 2.21 | 23.75 |
| T3 | 27.75 | 3.13 | 2.29 | 15.16 |

<sup>a</sup> Snapshots in which the tunnel is identified with respect to the total number of snapshots, in %.
<sup>b</sup> Data calculated with a probe radius of 2.0 Å.
<sup>c</sup> In Å.

To conclude, the active site of PlaF is connected to its surface with three tunnels. In the t-PlaF_A_ configuration, only T1 and T2 allow direct access of GPL or LGPL substrates from the membrane.

### PlaF preferentially hydrolyses medium-acyl chain LGPLs

Previously, we showed that PlaF can produce LGPLs by releasing FAs bound to the *sn*-1 position of GPLs. Here we experimentally tested if purified PlaF *in vitro* hydrolyses LGPLs by quantifying fatty acids released from a range of LGPLs varying in the head group and acyl chain length (C14-C18). The results revealed that PlaF can hydrolyze all tested LGPLs, with a preference for medium-acyl chain LGPLs (Figure 3). Interestingly, the lysoPLA activity of PlaF was 10- to 100-fold higher for all LGPLs than its PLA activity [19], indicating that hydrolysis of the first acyl chain in GPLs is much slower than that of the second one.

**Figure 3:**
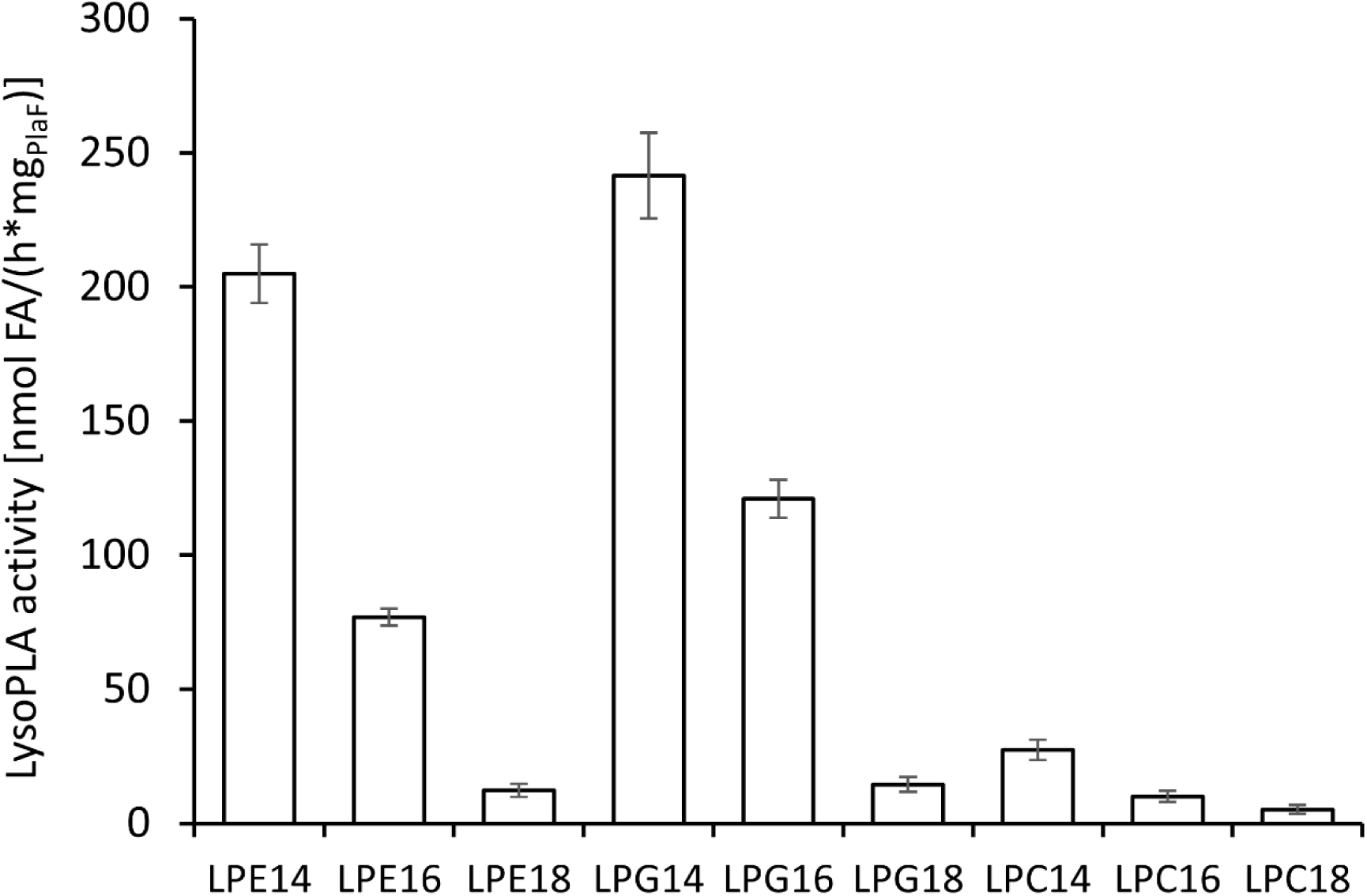
Lysophospholipase A activity of PlaF. Assays were performed by incubating purified PlaF (8 nmol) in *n*-dodecyl β-D-maltoside (DDM) micelles with the substrate, followed by quantification of released FA by NEFA-assay. Lyso-phosphatidylethanolamine (LPE), lyso-phosphatidylglycerol (LPG), and lyso-phosphatidylcholine (LPC) contain fatty acids with 14 - 18 carbon atoms. The results are the means ± standard deviation of three independent experiments.

### GPL and LGPL substrate extraction into solvent and acyl chain mobility

For probing the energetics of GPL and LGPL substrate extraction from the membrane into the solvent, we computed the free energy profile for DLPG and 1-myristoyl-2-hydroxy-*sn*-glycero-3-phosphoglycerol (2LMG) extraction (Supplementary results), which resulted in free energy differences between the two states of ∼13 ± 0.1 kcal mol^-1^ and ∼8 ± 0.3 kcal mol^-1^ (Figure S2A), in very good agreement with the excess chemical potential related to these lipids’ critical micelle concentration (CMC). For access to T3, substrates would need to leave the membrane and pass through the water phase, which makes this route energetically unfavorable. Hence, T3 was not considered for further analyses.

As T1 and T2 are immersed in the hydrophilic membrane surface (Figure S3A), access of GPL and LGPL substrates to the tunnels *via* the head groups is plausible. However, the tunnels’ diameters are much smaller than that of a GPL like DLPG while in the membrane (see above). To explore the possibility that lipids access via their acyl chain instead, we probed how frequently the terminus of a GPL’s acyl chain can reach the membrane interface. The probability distribution of GPL’s acyl chains with respect to the coordinate perpendicular to the membrane (z-coordinate) was determined during the last 40 ns of 300 or 100 ns long MD simulations for membrane bilayers with or without t-PlaF_A_, respectively (Figure S3A). Tails from both the upper and lower leaflet were considered. Positive *z*-coordinate values indicate that a tail moves towards the water-membrane interface of *its* leaflet; negative values indicate that it moves towards the interface of the *opposite* leaflet. The peak of the probability distributions is at *z* ≍ 2 Å indicating the mobility of lipid termini within the leaflet (Figure S3A, see also Movie S1 and Movie S2). The interface of the simulated membrane is at *z* ≍ 10 Å (Figure S3B). Notably, the cumulative probability of finding an acyl chain terminus at *z* > 10 Å is 1.5 % and 1.0 % for systems with or without PlaF, respectively. Hence, there is a finite likelihood that acyl chain termini can reach the entrances of T1 and T2. This result is also supported by the electron density profiles of the membrane components (Figure S3B).

To conclude, for t-PlaF, the access route of substrates to T3 is energetically unfavorable. By contrast, acyl chain termini of GPL lipids can reach the entrances of T1 and T2 during the time scales of our MD simulations.

### Access modes of GPL and LGPL substrates into PlaF

As a prerequisite to computing the energetics of substrate access to the active site of PlaF, we aimed to identify favorable access modes. We applied steered molecular dynamics (sMD) simulations [29] to pull substrates inside T1 and T2 (Figure 4) *via* head access first or tail access first.

**Figure 4:**
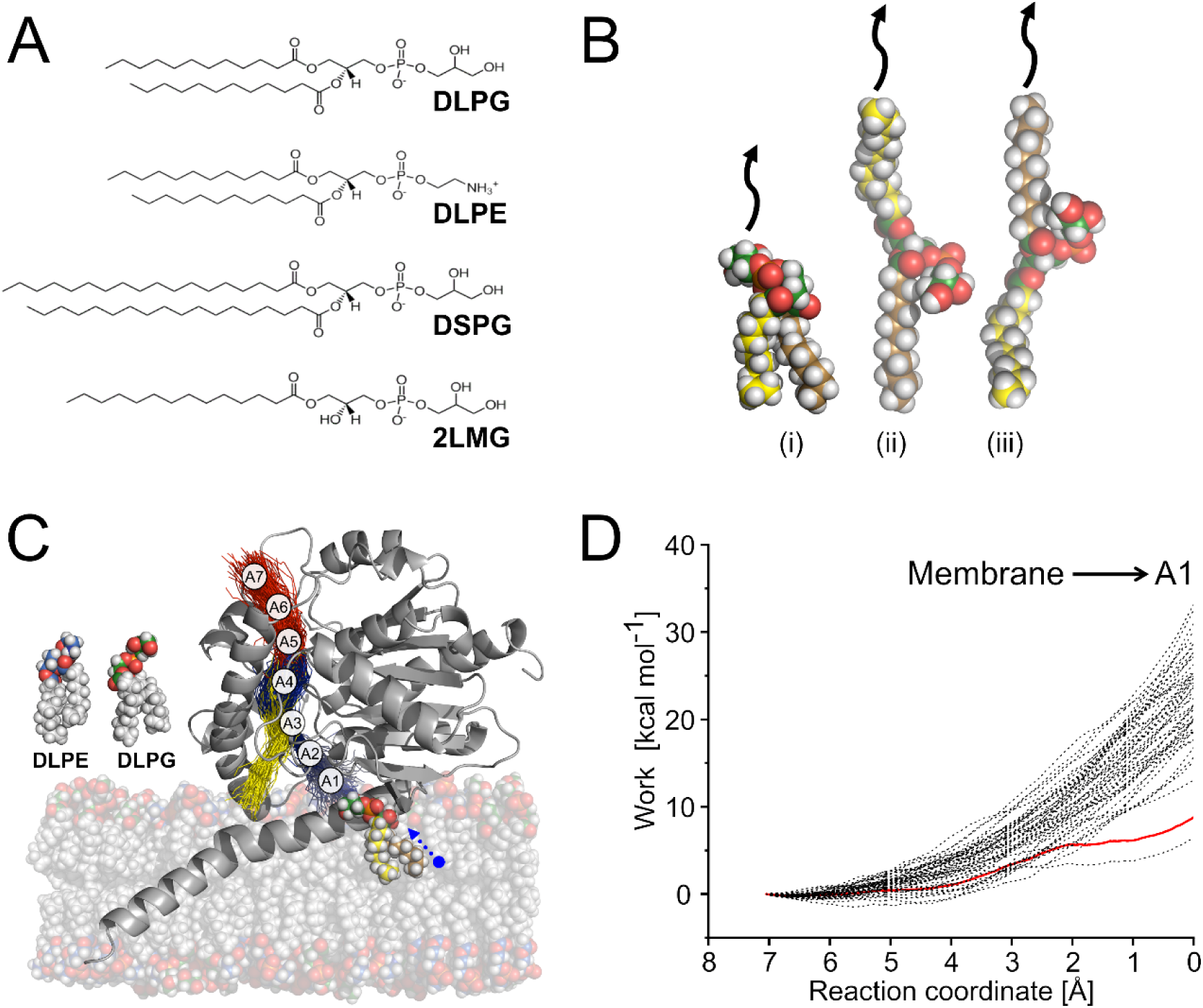
Illustration of the substrate access in t-PlaF_A_. A) Investigated GPL substrates, 1,2-dilauroyl-*sn*-glycero-3-phosphoglycerol (DLPG), 1,2-dilauroyl-*sn*-glycero-3-phosphorylethanolamine (DLPE), 1,2-distearoyl-*sn*-glycero-3-phosphoglycerol (DSPG), and LGPL substrate, 1-myristoyl-2-hydroxy-*sn*-glycero-3-phosphoglycerol (2LMG). B) Possible modes by which a GPL can access a tunnel (indicated with black arrow): with its head (green spheres represent the C atoms) first (i), tail 1 (yellow spheres represent the C atoms) first (ii), or tail 2 (orange spheres represent the C atoms) first (iii). C) PlaF is embedded in a membrane consisting of DLPE (head group C atoms as blue spheres) and DLPG (head group C atoms as green spheres) at a ratio of 3:1. The DLPG closest to the entrance of T1 (acyl chains colored) is shown while being loaded by its head, in the direction indicated with a blue arrow. A segmented path was considered for substrate access. T1 was segmented into four parts, and T3 into three parts, which are used as pulling points in sMD simulations. Depending on the access mode, in the last pulling step, the *sn*-1 or *sn*-2 site of the substrate is further pulled towards the nucleophilic OH group of the catalytic S137, resulting in, in total, 8 steps. A similar approach was used for T2 (Figure S4A). D) For the first segment of T1 (i.e., A1), the work done (black lines) during 50 independent replicas of sMD simulations to pull the DLPG from the membrane is plotted against the reaction coordinate. The coordinates of the replica, the work-versus-reaction coordinate profile of which is closest to the Jarzynski’s average (red line), are considered for pulling in the next segment. See the SI for plots of all other sMD simulations.

The closest substrate to the tunnel entrance was chosen for sMD simulations. The terminal oxygen and nitrogen atom of phosphatidylglycerol (PG) or phosphatidylethanolamine (PE) head groups, respectively, were considered for head access pulling. For tail access, the terminal carbons of respective acyl chains were considered. Substrates from the membrane were initially pulled through consecutive virtual points in T1 or T2 using four or five steps, respectively (Figure S4A, Table S2). However, pulling with terminal atoms leaves the cleavage site of the substrate distant to the catalytic S137 (Figure S4B). Therefore, the substrates were further pulled into T3, using three additional steps (Figure S4A). Depending on the access mode, the *sn*-1 or *sn*-2 sites of respective substrates were further pulled towards the nucleophilic OH group of the catalytic S137 (Table S2). Finally, this resulted in pulling pathways subdivided into eight and nine steps for T1 and T2, respectively (Table S2).

As a reaction coordinate, the distance between the pulled atom of a substrate and the consecutive virtual point was used. For each step, we repeated the pulling 50 times and computed the work done as a function of the reaction coordinate. By applying Jarzynski’s relation (eq. 1) [30], the work was related to the free energy difference between the two states of the pulling simulation. The sMD trajectory whose work-versus-reaction coordinate profile is closest to the Jarzynski average (eq. 1) was considered most favorable. Its endpoint provided the starting point for the sMD simulations in the next part of the pulling pathway. As a result, the access pathway is close to the lowest-free energy pathway of substrate access to the catalytic site. Overall, this approach is the reversed version of sampling unbinding trajectories of ligands from proteins before applying Jarzynski’s relation [31–33] but uses piecewise sMD simulations along the pathway to account for the curvilinear tunnels. A total of ∼27 µs of sMD simulation time was used for either tunnel (Table S3).

The activity of PlaF for GPL decreases with the increasing lengths of the acyl chain between C12 and C18, irrespective of the type of head group, PG or PE [19]. In addition, the number of acyl chains in a substrate also influences the PlaF activity, with LGPLs yielding a higher activity than GPLs (Figure 3). Hence, we chose DLPG with which PlaF is most active [19], 1,2-dilauroyl-*sn*-glycero-3-phosphorylethanolamine (DLPE), 1,2-distearoyl-*sn*-glycero-3-phosphoglycerol (DSPG), and 2LMG, a LGPL, for generating access modes (Figure 4A). Figure 4 exemplarily shows illustrations of the three access types for DLPG (see also Movies S3 - S8). Work-versus-reaction coordinate profiles for all pulling simulations related to DLPG access are shown in Figure S5 for T1 and Figure S6 for T2. Based on computed potentials of mean force (PMF) to evaluate the energetics of the access modes (see the next chapter), only tail access was considered for sMD simulations of the other GPL substrates (Figure S7). For 2LMG, head and tail access were considered for sMD simulations.

To conclude, seven access modes of GPL and two of LGPL substrates into PlaF were generated for T1 and T2, resulting in 18 access modes in total.

### Potentials of mean force of DLPG access modes

PMFs were computed from umbrella sampling (US) simulations [34] and post-processing with WHAM [35, 36] to evaluate the energetics of substrate access for the access modes described in the previous chapter (Figure 4). As a reaction coordinate, the distance between the center of mass (COM) of the three oxygen atoms of the glycerol moiety in the substrate to the COM of C_α_ atoms of the catalytic residues S137 and H286 was used. Residue D258 was not included in the reaction coordinate, as its side chain is distant from the active site (Figure S4A). As the tunnels are almost straight, the reaction coordinate monotonically decreases as the substrate approaches the active site from the membrane (Figure S4B). Initially, we focused on the US simulations for the best PlaF substrate [19], DLPG. PMFs were calculated for the three access modes of DLPG across either tunnel, T1 and T2. The PMFs were evaluated for convergence, excluding the first 200 ns of 300 ns sampling time. PMFs were found converged by 300 ns, yielding a maximal difference of ∼1 kcal mol^-1^ as to a PMF computed from 280 ns per window (Figure S8). The median overlap between the reaction coordinate distributions of neighboring windows was sufficient (≥ 4.8% and 3.5% for T1 and T2, respectively) (Figure S9).

The PMFs of DLPG access modes show marked differences (Figure 5A). Access with the head first is the least favorable for both T1 and T2, resulting in steep PMFs with free energy barriers of 11 and 9 kcal mol^-1^ (Figure 5A), in contrast to tail access. Most of the residues within a radius of 3 Å in T1 and T2 have either a neutral non-polar side chain, which likely facilitates tail access to the active site of PlaF. Furthermore, access with either one of the two tails first is more favorable in T2 than T1 (Figure 5A). Finally, access with tail 1 first in T2 is most favorable and results in no free energy barrier until the substrate reaches the active site (Figure 5A). As the two acyl chains of DLPG are identical, these results suggest that their connection with the glycerol moiety causes differences in how the lipid interacts with the tunnel, which may explain how PlaF achieves regioselectivity to exert its PLA_1_ function.

**Figure 5:**
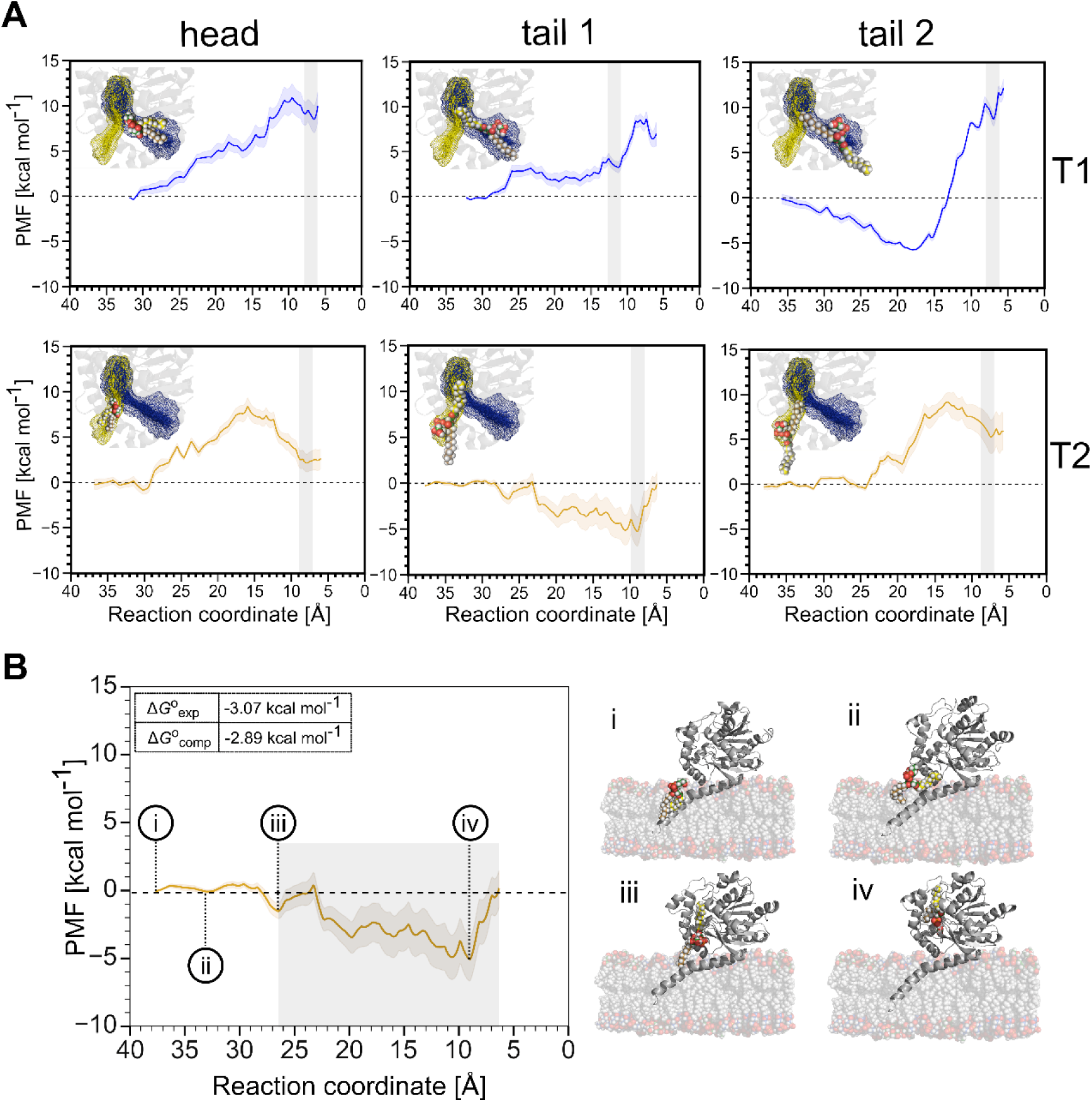
Potential of mean force profiles for DLPG access. (A) PMFs of three access modes (head, tail 1, tail 2; see Figure 4B) of DLPG in T1 (blue curve) and T2 (yellow curve). For both tunnels, access with tail 1 first yields the lowest free energy barriers to reach the active site. Furthermore, DLPG access into T2 with tail 1 first is overall the most favorable. The catalytic site is marked with a grey box. Insets within the plots illustrate the different DLPG access modes into the respective tunnels. (B) States during DLPG access via tail 1 through T2, shown on the right, are marked in the PMF profile (left). The grey box corresponds to the integration limits used to calculate *K*_eq_ (eq. 2) to determine 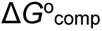 (see inset). State i: The starting position of DLPG (in the membrane). State ii: Tail 1 reaches the surface of the membrane close to the entrance of T2. State iii: Tail 1 enters inside T2, while tail 2 remains within the membrane. State iv: *sn*-1 site of tail 1 reaches the catalytic site.

To validate our results, we computed the absolute binding free energy of DLPG to PlaF from the PMF for tail 1 access in T2, 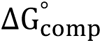 = −2.89 ± 1.46 kcal mol^-1^ (eq. 4). Assuming that product formation is slower than substrate dissociation from an enzyme, the Michaelis constant *K*_m_ is equal to the dissociation constant *K*_D_ of the enzyme-substrate complex [37, 38]. Under this assumption, from *K*_m_ = 7.612 ± 1.907 mM for DLPG in PlaF [39], the experimental binding free energy 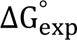 = −3.07 ± 0.30 kcal mol^-1^ at *T* = 303 K is calculated, which is within chemical accuracy [40] of 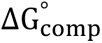.

We also computed 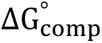 for the other five access modes of DLPG (eq. 4). The lowest 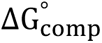 among all six modes was obtained for tail 2 access in T1 (Table S4). However, the PMF profile (Figure 5A) reveals that the configurational free energy minimum is not situated close to the active site but in the middle of T1. Here, one of the tails is still in the membrane, while the other is being loaded into the tunnel. If the PMF profile is integrated with two separate parts, first, a negative free energy for tail access into the tunnel results, followed by a positive free energy to reach the active site. This suggests that this access mode cannot yield a catalytically active configuration. For the other four access modes, 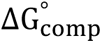 > 0.96 kcal mol^-1^ (Table S4). These findings corroborate tail 1 access of DLPG in T2 as the most likely access mode.

Along the PMF of tail 1 access of DLPG in T2, four distinct states can be identified (Figure 5B). The two tails of DLPG are immersed in the membrane at a reaction coordinate value of ∼38 Å from the active site (Figure 5B, state i). The PMF remains essentially unchanged if tail 1 approaches the surface of the membrane and the entrance of T2 (state ii). This is concordant with the tail distributions along the z-coordinate during unbiased MD simulations (Figure S3), indicating that tail termini can reach one of the access tunnels of t-PlaF_A_ without a considerable energetic cost. Once tail 1 enters T2, the PMF becomes negative (state iii), indicating that DLPG access that way is favorable. Finally, at ∼8 Å of the reaction coordinate, the PMF has a global minimum (state iv). There, tail 1 is located in T3, and the acyl moiety at the *sn*-1 position of DLPG is close to the catalytic S137 of PlaF (Figure S10B) such that a nucleophilic attack can commence.

To conclude, we identified T2 as the preferred access tunnel for DLPG in PlaF. Access with tail 1 first is most favorable there. This is in line with PlaF being a PLA_1_, which cleaves its substrates at the *sn*-1 position. As of T3, it is likely essential for substrate access by allowing to accommodate the substrate tail to be hydrolyzed by PlaF.

### Potentials of mean force for accesses of other substrates

Considering the results for DLPG, we performed US simulations for DSPG and DLPE only for tail 1 access. For the LGPL substrate, it has remained undetermined if the head or tail access is energetically favorable; hence, we performed US simulations for both access modes of 2LMG. As for DLPG, T2 is preferred over T1, regardless of the access modes (Figure 5A), we only considered T2 for computing PMFs for the other substrates. Similar to DLPG, the PMFs converged at 300 ns of sampling time, yielding a maximal difference of ∼0.5 kcal mol^-1^ as to a PMF computed from 280 ns per window (Figure S11). Neighboring umbrella windows have a sufficient median overlap ≥ 3.2% (Figure S12).

For DSPG and DLPE, access with tail 1 first in T2 results in pronounced free energy barriers of 11 and 14 kcal mol^-1^ (Figure 6A, B), in contrast to DLPG (0.5 kcal mol^-1^). This finding indicates that a longer acyl chain or a neutral head group makes substrate access to PlaF disfavorable, which coincides with lower PlaF activities for such substrates [19]. For 2LMG, access with the tail first is more favorable than with the head, as for DLPG (Figure 6C, D). Furthermore, tail access by 2LMG leads to a free energy barrier lower by ∼6.5 kcal mol^-1^ than those for tail access by DSPG and DLPE (Figure 6A, B, D), which is concordant with the activity profile of PlaF [19].

**Figure 6:**
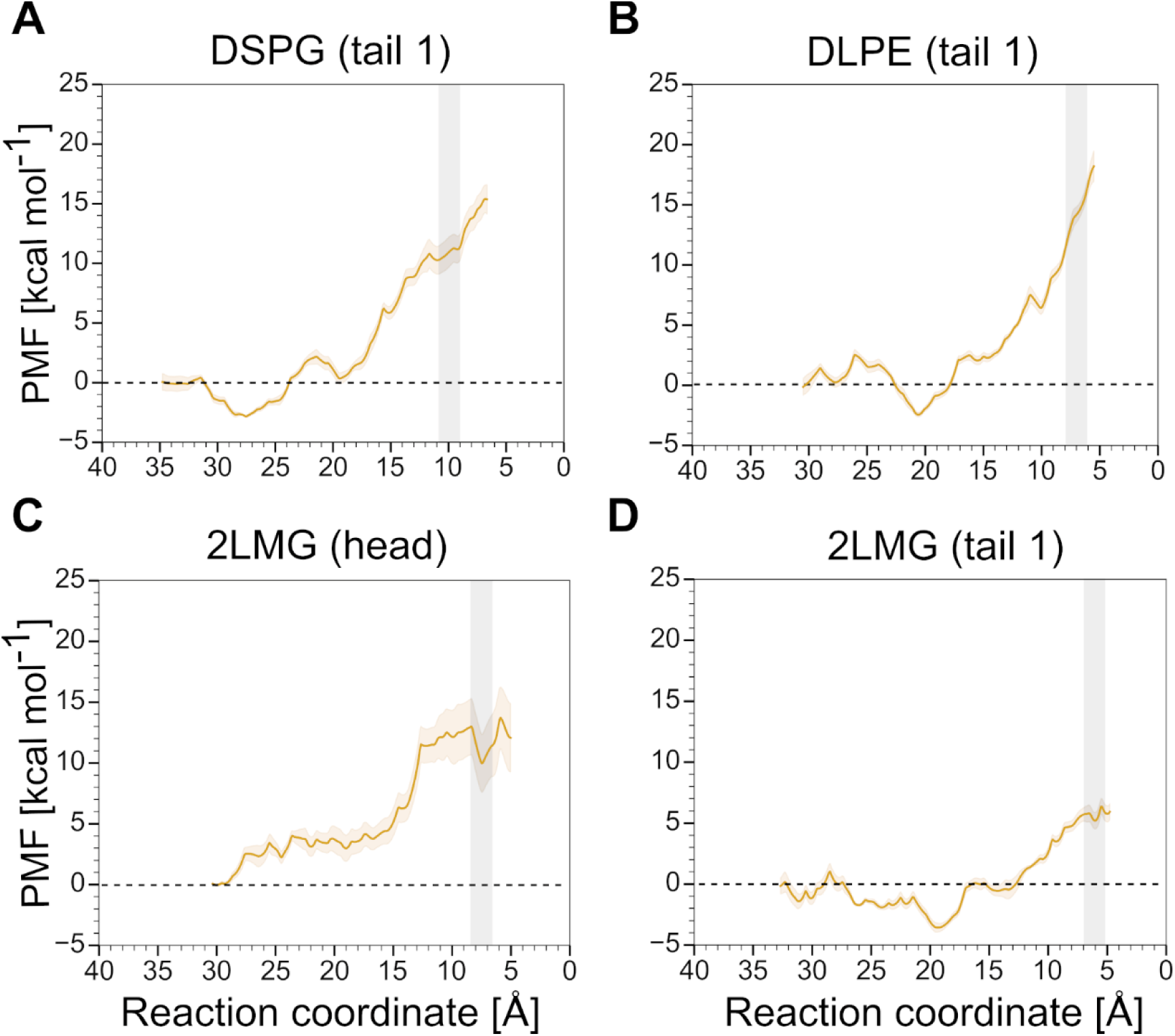
PMF profiles for other substrates across T2. Four systems were investigated to reveal the energetics of DSPG access via tail 1 (A), DLPE via tail 1 (B), 2LMG via head (C), and 2LMG via tail 1 (D). Among these substrates, access of 2LMG via tail 1 has the lowest free energy barrier. The catalytic site is marked with a grey box.

To conclude, tail 1 access in T2 of GPL substrates with longer acyl chains or neutral head groups is disfavorable compared to DLPG access, in line with PlaF’s substrate specificity. For the LGPL substrate 2LMG, tail 1 access is also favored over head access and more favorable than DSPG and DLPE access.

### Tryptophan substitutions in T2 hamper DLPG access

To validate the prediction that T2 is the preferred access pathway, we identified residue positions in all identified tunnels that, when substituted with tryptophan (Trp), should constrict the tunnel and, thus, block substrate access. Earlier, this strategy has been used to block tunnels of a dehalogenase and influence its activity by limiting the rate of product release [41]. In the case of PlaF, the products are less bulky than the substrates, such that product release should be less impacted than substrate access due to constricted tunnels.

PlaF variants were predicted subject to minimizing the structural destabilization due to the Trp substitution and preferring sites within the tunnels that influence its geometric characteristics (Table S5). We predicted four Trp substitutions for T1 and five for T2, and T3 each (Table S5). With any one of these substitutions in place, the impacted tunnel could not be identified anymore by CAVER applying the previously used probe radius of 2 Å, but with a smaller probe radius of 1.2 Å (Figure 7). This indicates their constriction, also displayed by the time evolution of the tunnel profiles of the PlaF variants compared to PlaF wild type (Figure S13).

**Figure 7:**
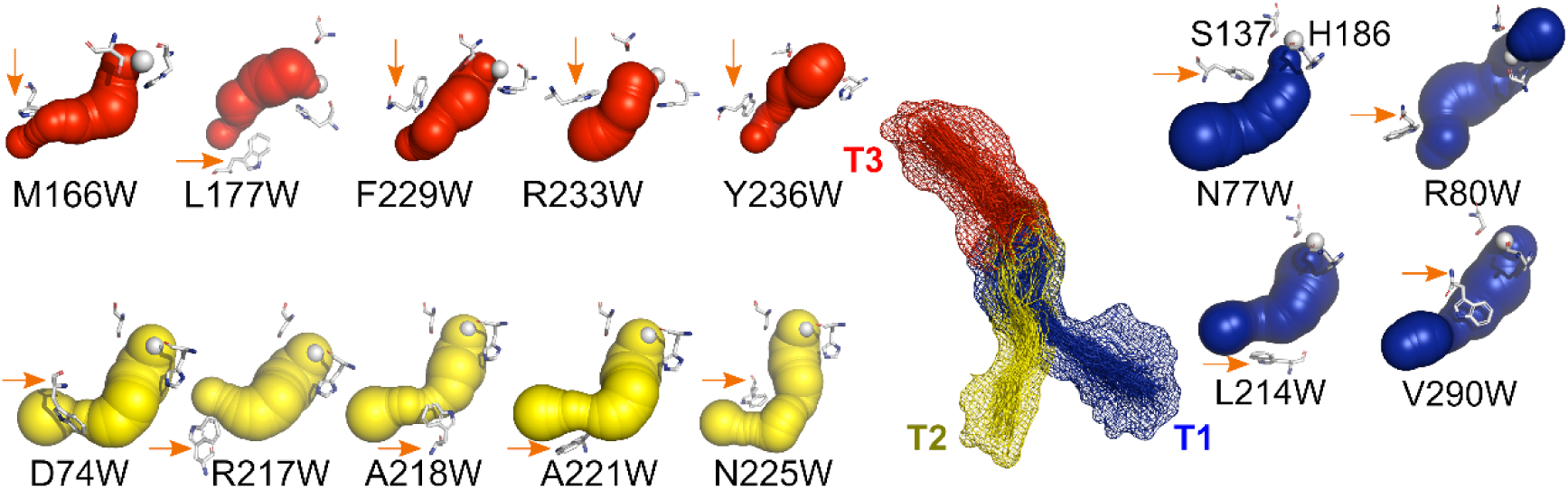
Influence of tryptophan substitutions on the radius of PlaF tunnels. The tunnels T1 (blue), T2 (yellow), and T3 (red) are identified by CAVER with a reduced probe radius of 1.2 Å, instead of 2 Å used otherwise (Figure 2), showing that the tryptophan substitutions (orange arrows) narrow the tunnels. White spheres, wherever visible, represent the origin of the search defined by the COM of the catalytic residues S137 and H286.

The mutations of fourteen suggested residues (Figure 7) to Trp were generated by sequence- and ligation-independent cloning (SLIC) method in which the whole *p*-PlaF expression vector was amplified. Mutations were verified by sequencing, and the wild type PlaF (PlaF_WT_) and respective variants were produced in the homologous host, *P. aeruginosa,* following their immobilized metal affinity chromatographic (IMAC) purification from membranes solubilized with *n*-dodecyl β-D-maltoside (DDM) (Figure S14). All variants showed purity comparable to that of PlaF_WT_ (Figure S14). The specific activity of PlaF variants and PlaF_WT_ was analyzed by measuring the hydrolysis of small (*p*-nitrophenyl butyrate, *p*-NPB) and large (DLPG) substrates (Figure 8A). The activities of all nine variants in T1 and T3 measured with *p*-NPB and DLPG were similar to the activity of PlaF_WT_. In contrast, all five T2 variants had a significantly (p < 0.001) lower activity with *p*-NPB and DLPG than PlaF_WT_.

**Figure 8:**
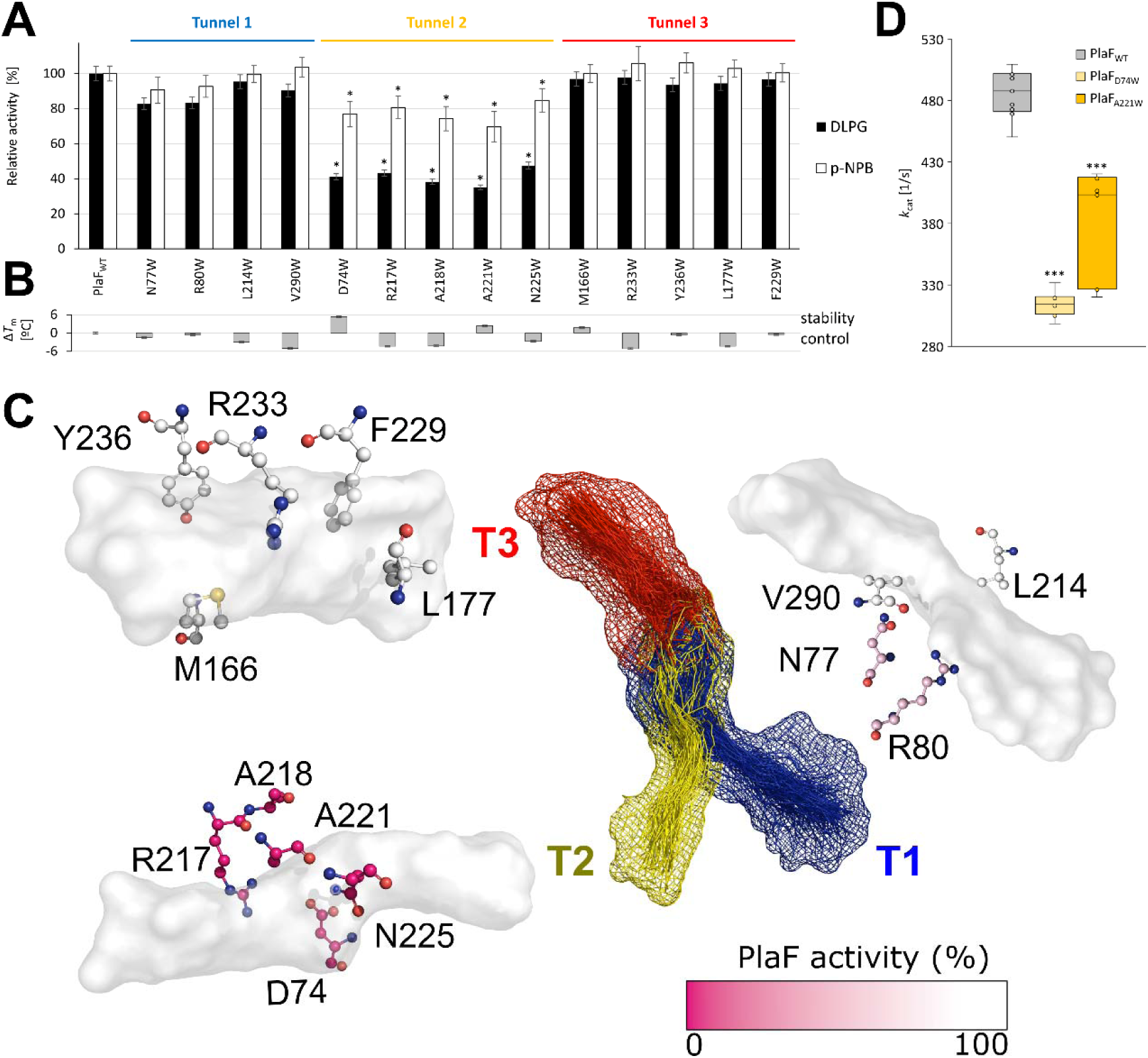
Lipolytic activity of PlaF and variants with Trp substitutions in T1-T3. A) Enzyme activities of purified PlaF_WT_ and variants carrying respective substitutions measured with DLPG and *p*-NPB. Activities are normalized to the activity of PlaF_WT_, which was set as 100%. Results are means ± standard deviation of three independent measurements. Statistical analysis was performed using the t-test (* *p* < 0.001) of normally distributed values for DLPG (*n* = 8) and *p*-NPB (*n* = 9) measurements. B) The thermal stabilities of purified PlaF_WT_ and variants were measured by nanoDSF. Results are shown as a difference in the melting temperatures (Δ*T*_m_) of the respective PlaF variant and PlaF_WT_, which was 57.3 ± 0.2 °C. Results are means ± standard deviation of three independent measurements, each performed with three samples. C) The tunnels, T1 - T3 (mesh view in the center) are represented as white surfaces. The investigated amino acids are shown in ball-and-stick representation. The CPK coloring scheme was used to color all atoms except carbons, which vary from pink to white (see color scale) related to the PlaF activity for DLPG after substituting the corresponding residue for a tryptophan. The PlaF activity is reduced the most if Trp substitutions involve T2. D) Kinetic parameters of PlaF_WT_ and the substrate binding-T2 variants PlaF_D74W_ and PlaF_A221W_ measured using *p*-NPB. Kinetic parameters were determined by non-linear regression analysis of the data (*n* = 9 for PlaF_WT_ and PlaF_A221W_, *n* = 6 for PlaF_D74W_) fitted to the Michaelis-Menten equation. The box plots represent the interquartile range between the first and third quartiles of the kinetic parameters determined for PlaF_WT_, PlaF_D74W_, and PlaF_A221W_. The line inside the box is the median, and the whiskers represent the lowest and highest values. Statistical analysis was performed using the t-test (*** *p* < 10^-5^).

To exclude that a substitution leads to an unstable protein, we measured the thermal stability of each variant by detecting intrinsic protein fluorescence upon unfolding. None of the variants showed a drastically reduced stability (Figure 8B). On the other hand, two T2 variants were slightly more stable (1.8 - 2.4°C) than the PlaF_WT_, and three variants were slightly less stable (2.7 - 4.3°C). Hence, Trp mutations do not affect PlaF’s stability at the temperature of enzymatic assays (30°C).

The observation that PlaF activities with DLPG and *p*-NPB predominantly decreased with substitutions in T2 (Figure 8A) indicates that substitutions with the bulky Trp impact passage through T2 (Figure 8C). As expected, the activities with the larger DLPG decreased more (52 - 65%) than with the smaller *p*-NPB (15 - 30%).

We also determined kinetic parameters for the *p*-NPB hydrolysis of PlaF_A221W_ and PlaF_D74W_ with substitutions in T2 (Figure S15). Despite a less prominent effect of the substitutions on specific activities measured with *p*-NPB than DLPG, *p*-NPB allows for reliable determination of PlaF activities over the range of substrate concentration and, thus, is applicable for the determination of kinetic parameters of PlaF. In contrast, enzyme kinetic experiments using hydrophobic DLPG are not feasible because of micelle formation and the slow rate of reaction.

Although, the affinities of PlaF_WT_, PlaF_A221W_, and PlaF_D74W_ for *p*-NPB are similar (Figure S15, also see the table in the inset), the catalytic turnover of both variants (PlaF_A221W_: *k*_cat_ = 314.6 ± 7.0 s^-1^; PlaF_D74W_: *k*_cat_ = 403.4 ± 15.1 s^-1^) was significantly (*p* < 10^-5^) lower than of PlaF_WT_ (*k*_cat_ = 487.8 ± 15.4 s^-1^) (Figure 8D). These results confirm that the point mutations PlaF_A221W_ and PlaF_D74W_ interfere with *p*-NPB access through T2.

To conclude, biochemical studies of fourteen PlaF variants with Trp substitutions introduced in all three tunnels showed tha only substitutions in T2 reduced lipolytic activity of PlaF. These results confirm that T2 is the main route for substrate access from the membrane to the catalytic site.

### Potential egress pathways of PlaF products

Next, we aimed at identifying potential egress pathways for products of PlaF-catalyzed hydrolysis. We performed a set of unbiased MD simulations starting from a hydrolyzed 2LMG in t-PlaF_A_. The starting coordinates were taken from the last snapshot of the US simulations of 2LMG with tail 1 access through T2, considering the umbrella window where the *sn*-1 position of 2LMG was closest to the catalytic site. Then, 2LMG was cleaved into the respective products without changing their orientation in the tunnels (Figure 9A). This led to MYR being in T3 at the beginning of the simulations and the PGR (phosphatidylglycerol from LGPL, 2LMG) moiety pointing towards T2 (Figure 9B).

**Figure 9:**
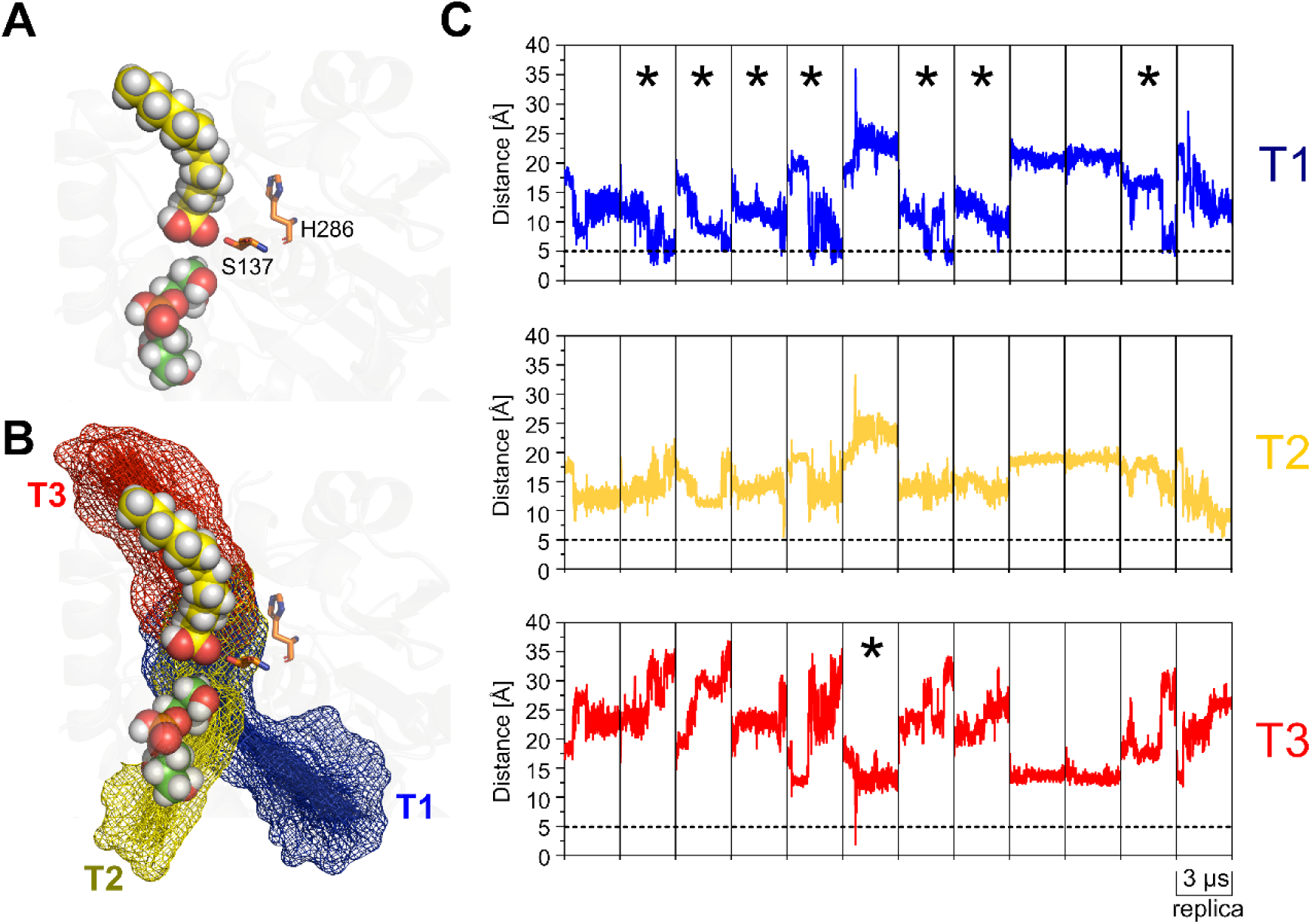
Unbiased MD simulations of t-PlaF_A_ with bound hydrolysis products. (A) Starting configuration of the 2LMG products in t-PlaF_A_. MYR is represented with yellow spheres and PGR with green spheres. The catalytic S137 and H286 are shown as orange sticks. (B) The products are mapped over the respective tunnels. (C) The distance of MYR to the entrances of T1-T3 during 12 replicas of unbiased MD simulations of 3 µs. The dashed black line depicts the chosen cutoff of 5 Å, with replicas that reach this cutoff marked with an asterisk. MYR reaches a distance ≤ 5 Å to the entrance of T1 in 7 replicas, in 1 replica for T3, and in none for T2. Note that in chain A of the PlaF crystal structure, MYR is found in T1.

In 12 replicas of 3 µs length each, the products relocated within the tunnels, sometimes even diffusing into the solvent (PGR moiety in 2/12 replicas via T1 and 3/12 replicas via T2; Figure S16). MYR relocated from its original position in T3 and approached the other tunnels of PlaF during the course of the MD simulations (Figure 9C). To deduce the displacement of MYR, we measured the distance of the carboxyl carbon to the entrance of each tunnel. A cutoff of 5 Å, according to previous studies [42–44], was used to identify those replicas where MYR reaches close to the tunnel entrance. MYR moved in 7/12 replicas to the entrance of T1 and in 1/12 replicas to the entrance of T3; the entrance of T2 was not reached (Figure 9C). Interestingly, the instance of MYR reaching the entrance of T3 flips within T3 such that the carboxyl group points to the entrance, rather than to the active site as after hydrolysis (Figure 9B). Altogether, MYR reaches the entrance of T1 significantly more frequently than T2 (*p* = 0.0008) and T3 (*p* = 0.0047) (Figure S17, Table S6).

To conclude, hydrolysis products of 2LMG diffuse within PlaF during time scales of 3 μs, sometimes also between tunnels. T1 and, to a lower extent, T3 are the most likely egress pathways of FAs from PlaF, although more sampling is required to observe actual egress.

## Discussion

Dimer-monomer transitions regulate the activity of several membrane-bound phospholipases, including PLA_1_, and PLA_2_ [45–52]. Previously, we showed that PlaF becomes active due to a dimer-to-monomer transition followed by tilting of the monomer in the membrane, resulting in t-PlaF_A_ being the active configuration of PlaF [19]. Here, we addressed the questions of how membrane-bound substrates reach the active site of PlaF_A_ and how the characteristics of the active site tunnels determine the activity, specificity, and regioselectivity of PlaF for medium-chain substrates. We performed unbiased and biased MD simulations and showed by configurational free energy computations and mutational and enzymatic studies for t-PlaF_A_ that A) access of the two main PlaF substrates DLPG and 2LMG occurs most likely through tunnel T2 in a tail-first mode, B) access of substrates with longer acyl chains or neutral head groups is less favorable, C) tail 1 access of DLPG and 2MLG in T2 is more favorable than tail 2 access, D) T3 accommodates the substrate tail to be hydrolyzed, and E) T1 and T3 are potential product egress pathways.

Previous studies indicated that the characteristics of substrate access tunnels can have a decisive influence on enzyme-substrate specificity and activity [53–56]. In t-PlaF, we focused on T1 and T2 because only these two allow direct access of GPL or LGPL substrates from the membrane in the t-PlaF_A_ configuration. By contrast, to enter into T3, substrates would need to pass through the solvent, which is energetically unfavorable. In di-PlaF, T2 is closest to the membrane with a distance of 7.4 ± 1.5 Å but T1 and T3 are at a distance > 12 Å (Figure S18A). Hence, we also investigated substrate access to T2 in di-PlaF.

For assessing the energetics of substrate access, first, we generated 18 pathways, considering GPL and LGPL as substrates in T1 and T2 using sMD simulations. By relating the work along the reaction coordinate to the free energy difference between two states of the pulling simulations via Jarzynski’s relation and considering the endpoint of the sMD trajectory closest to the Jarzynski average as the starting point for the next sMD simulation, we obtained low-free energy pathways of substrate access to the catalytic site. sMD simulations have been widely used to explore similar biological processes such as the loading of GPL substrates into human phospholipase A_2_ (PLA_2_) [57] or recognition of arachidonic acid by cytochrome P450 2E1 across the access channel [58]. The pathways served for defining reference points for subsequent US simulations, such that distributions of sampled states sufficiently overlapped, which is essential to yield accurate results in PMF computations [59]. Applying US along pathways identified by sMD simulations [60] or targeted simulations [61, 62] has been shown to be an effective method of computing PMF. Moreover, the choice of an appropriate reaction coordinate is essential for this approach [63–65]. We probed for the convergence of our PMFs by comparing PMFs generated from increasing lengths of US simulations and found that US times of ∼300 ns are needed to yield PMF differences below chemical accuracy [66]. Finally, we validated our PMF computations by comparing the computed absolute binding free energy of DLPG to PlaF for the most preferred access mode to an estimate of the experimental binding free energy.

The PMFs revealed that tail-first access through T2 is most preferred for DLPG and 2LMG. This finding is in line with the geometric analysis of T2, which revealed a tunnel bottleneck radius about half as large as the radius of DLPG deduced from the lipid’s area-per-lipid, which can explain why a headgroup-first access is disfavorable for steric reasons. Furthermore, we showed that acyl chains of lipids embedded in a membrane can reach the interface region in unbiased MD simulations and, thus, can interact with the tunnel entrance. Such protrusions of lipid tails occur on a timescale of approximately 100 ns depending on the extent of solvent exposure [67]. Tail-first access of GPLs into the active site has also been found for cyclopropane fatty acid synthase [68]. Tail-first access through T2 is favored because of the predominant hydrophobic nature of the tunnel walls. By contrast, T1 contains a higher number of charged Asp and Arg residues and fewer neutral residues than T2, which makes tail-first access there less favorable. In particular, the side chain of R80 protrudes into T1 at the tunnel entrance, which is reflected in an energy barrier of ∼3 kcal mol^-1^ found there for tail-first access.

Modifications in tunnels that connect a buried active site to the bulk solvent have been shown to affect ligand binding and unbinding [41]. Tunnel residues situated away from the active site are suitable targets for mutagenesis, as their replacement should not lead to a loss of the functionality of the active site [69]. Considering this, we introduced Trp substitutions to each of the three tunnels of PlaF and measured the activity of these PlaF variants. The Trp substitutions decreased PlaF’s lipolytic activity for small and large substrates only when introduced in T2, which suggests that T2 is involved in substrate access. However, from such steady-state experiments, it cannot be excluded that the Trp substitutions influence product egress, too [54].

Among the investigated substrates, higher energy barriers for access to the active site were found for those with longer acyl chains and neutral head groups, concordant with PlaF’s activity profile [19]. This finding may be explained with differences in the energetics of GPL self-assembly, which is influenced by the hydrocarbon chain length and the polarity of the head group [70]: Longer hydrocarbon chains and less polar head groups foster self-assembly, which would lead to higher energy barriers for leaving this equilibrium state [71] and entering into PlaF. These results indicate that the energetics of access of a membrane GPL substrate to the active site through tunnel T2 contributes to the substrate specificity of PlaF.

Furthermore, of the two constitutopic acyl chains in DLPG, access via tail 1 in T2 is energetically preferred over tail 2 access. If tail 1 enters first, the carbonyl oxy group at C1 of the glycero moiety can come closer to the nucleophilic S137 than if tail 2 enters first, (Figure S10) leading to preferential hydrolysis of the carboxylic ester bond at C1. Likewise, the regioselectivity of human 5-lipoxygenase is determined by the head/tail-first type orientation of its main substrate arachidonic acid in the active site [72]: The arachidonic acid can be positioned in the holoenzyme active site with both head-first and tail-first orientation, but only the tail-first orientation results in a configuration that yields 5-lipoxygenating activity. These results indicate that the tail-first access mode of a diacyl GPL substrate determines the regioselectivity of PlaF for hydrolysis of the acyl chain bound to the *sn*-1 position.

As T3 is oriented to the membrane neither in the monomeric nor in the di-PlaF configuration, it likely does not contribute to substrate access. We suggest the role of T3 to accommodate the acyl chain of substrates before and products after hydrolysis. T3, with a length of ∼15 Å, provides adequate space for substrates with medium-lengths of acyl chains and, thus, may affect the specificity of PlaF. Substrate tunnels that accommodate acyl chains hydrolyzed from their respective precursors have also been described for cholesterol acyltransferases [73]. Likewise, lipid phosphate phosphatases harbor such a cavity, accommodating the substrate’s acyl chain for optimal catalysis [74]. Site-directed mutagenesis in *Candida rugosa* lipase 1 revealed the role of such tunnels in determining the acyl chain length specificity [75].

As to di-PlaF, tail 1 access of DLPG across T2 revealed a free energy barrier of ∼13 kcal mol^-1^ (Figure S18B), in contrast to no free energy barrier in t-PlaF_A_ (Figure 5B). This high barrier may arise because of the location of T2 in di-PlaF, ∼7 Å above the membrane. Thus, substrates would need to pass through the solvent to enter T2. These findings indicate that di-PlaF is catalytically inactive, as determined experimentally [19], because of energetically unfavorable substrate access.

Our results from unbiased MD simulations of products suggest that T1 and, to a lower extent, T3 are egress pathways of FAs. As to T1, this suggestion is in agreement with the crystal structure of PlaF, where FAs are found in T1 [19]. In the tilted orientation of PlaF, FAs egressing via T1 would interact with the membrane interface and could diffuse into it. FAs in a membrane can affect its fluidity and permeability and protein-lipid interactions, thereby regulating important cell processes including signal transduction, motility, and biofilm formation [76, 77]. Via T3, they would egress into the periplasmic space. Anchored to the cytoplasmic membrane, PlaF is not a toxin targeting the host cell membrane but it has a direct influence on virulence adaptation of *P. aeruginosa* by modulating the membrane GPL composition [19]. However, it is unknown if FAs released from GPLs by PlaF are targeted to the external environment as for example diffusible FAs involved in cell-to-cell signaling [17, 18]. In this case, egress of FAs *via* T3 to the periplasm and their further passive diffusion or active transport would be possible [78].

In summary, we identified T2 as the preferred tunnel for substrate access to t-PlaF_A_, while T1, and to a lesser extent T3, are likely egress routes for FAs. The energetically favorable tail 1 access of substrates is in agreement with PlaF’s PLA_1_ function. The higher preference of PlaF for GPLs with medium-length acyl chains may be due to differences in the energetics of self-assembly and the length of T3, which accommodates them for hydrolysis (Figure 10). Finally, while t-PlaF_A_ enables substrate access to the active site, substrate access to di-PlaF is energetically unfavorable. Our results provide an atomistic-level understanding of the unique structural feature of PlaF that its function is dependent on monomerization followed by global reorientation of the single-pass TM protein at the membrane. They may furthermore aid in understanding the feedback regulation of PlaF, which is inhibited by FAs, and open up opportunities for developing potential drugs that inhibit PlaF to combat *P. aeruginosa* virulence during infections.

**Figure 10:**
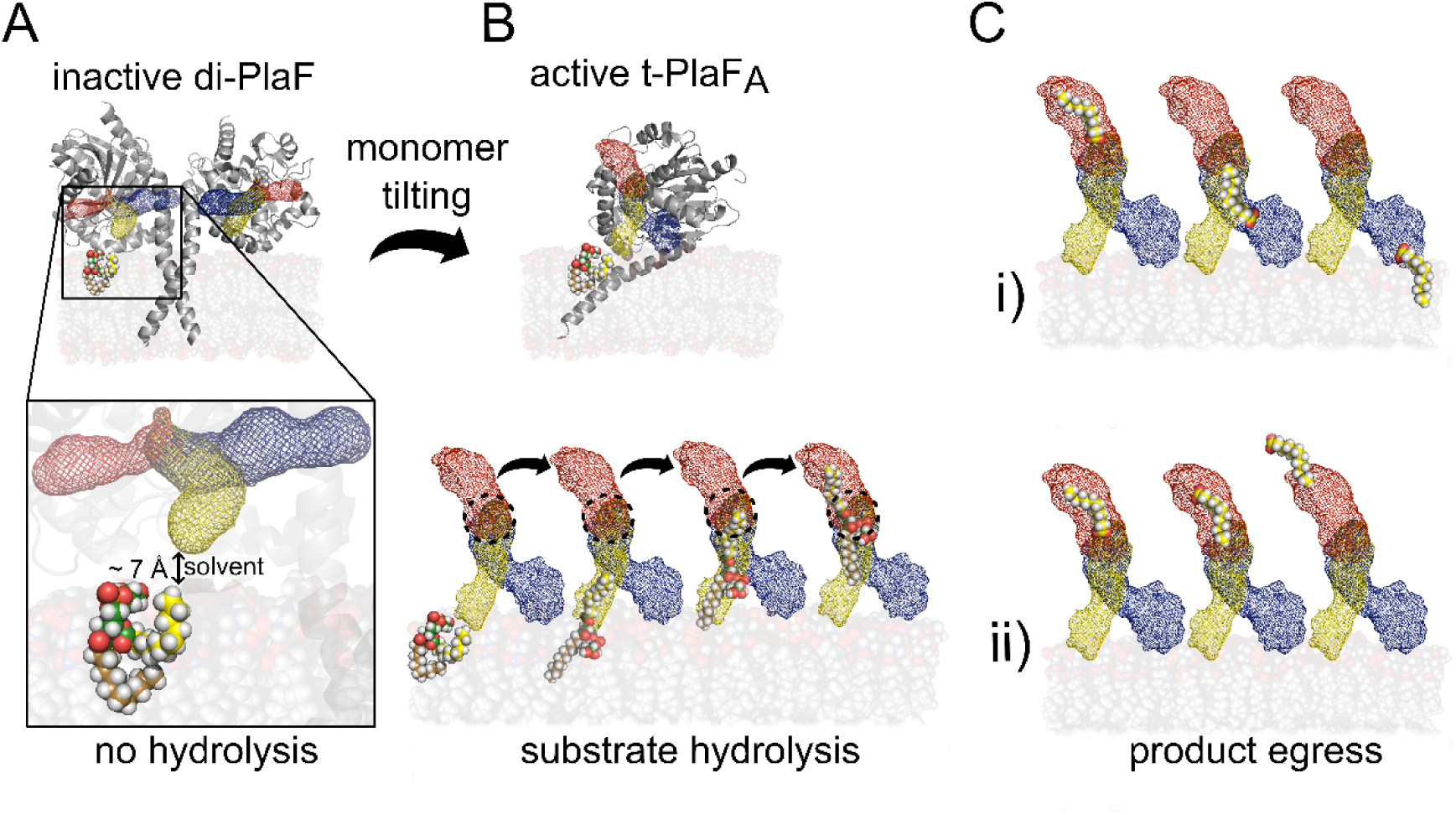
Schematic model of the mechanism of PlaF activity regulation. A) A higher concentration of PlaF results in the formation of an inactive dimer [19]. In di-PlaF, T2 is closer to the membrane than the other tunnels, but, still, the distance from the membrane interface is ∼7 Å, which requires the substrate to pass through the solvent. Hence, this configuration leads to inactive PlaF. B) At low concentrations, monomeric PlaF shows PLA_1_ activity and adopts a tilted configuration [19]. In the tilted configuration, PlaF orients such that T1 and T2 come close to the membrane interface. Substrate access occurs *via* tail 1 into T2. The acyl chain reaches the active site (dashed black circle) of PlaF, but the *sn*-1 cleavage site is still away from the active site. Further loading of the acyl chain requires it to enter into T3, and the substrate is hydrolyzed. C) After hydrolysis, the FA is in T3. i) Now, either FA relocates into T1, with the carboxyl group towards the entrance. At the T1 entrance, FA can interact with the membrane interface and diffuse into it. ii) Alternatively, the FA can flip around, such that the carboxyl group faces the T3 entrance, from where it can exit into the periplasmic space.

## Materials and methods

### Identification of the access tunnels

Tunnels emerging from the active site of PlaF were identified using CAVER 3.0 [26]. The COM of the catalytic residues S137 and H286 was defined as the starting point of the search, from which the possible connections of the tunnels to the bulk solvent were identified. The catalytic residue D258 was not included in this search criteria since its side chain is distant from the catalytic site. Probe and shell radii of 2 Å and 6 Å were used, respectively. The probe radius of 2 Å is slightly larger than the van der Waals radius of a phosphorous atom (i.e., 1.8 Å), present in every PlaF substrate to be investigated.

### Starting structure preparation

The crystal structure of PlaF is available from the Protein Data Bank (PDB) [79] (PDB id: 6I8W) [19]. The first five residues of the C-terminus were missing in the structure and, hence, were added using MODELLER [81]. The starting configuration of PlaF for MD simulations was prepared by embedding t-PlaF_A_ into a lipid bilayer membrane consisting of 75% DLPE and 25% DLPG. The tilted configuration of PlaF embedded in the membrane was predicted by the Positioning of Proteins in Membrane (PPM) method [82]. The head group composition of the membrane closely resembles that of the inner membrane of Gram-negative bacteria [8, 83, 84]. The prepared structure was used to investigate the loading mechanism of DLPG or DLPE into t-PlaF_A_. Furthermore, loading of DSPG and an LGPL, 2LMG, were also investigated. For that, t-PlaF_A_ was embedded into a membrane consisting of ∼10% of DSPG and 2LMG in the upper leaflet. The GPL composition in the lower leaflet of these systems is the same as that used for investigating DLPG and DLPE. The systems were prepared and solvated using CHARMM/GUI [85] or PACKMOL-Memgen [86]. A distance of at least 15 Å between the protein or membrane and the solvent box boundaries was used. To obtain a neutral system, counter ions were added that replaced solvent molecules. The size of the resulting systems was ∼140,000 atoms.

Systems excluding the t-PlaF_A_, but including one of the GPL substrates (i.e., DLPG) and one of the LGPL substrates (i.e., 2LMG), were also prepared to compare and decipher the energetics of lipid extraction from the membrane into solvent. Considering the orientation and position of t-PlaF_A_ in the membrane, one can safely assume that only substrates located in one leaflet will contact the catalytic domain of t-PlaF_A_ and, hence, have direct access. Therefore, the composition of one leaflet was slightly modified to reflect the inclusion of the selected substrate. For this, a ratio in the upper leaflet of 6:2:1 for DLPE, DLPG, and the respective substrate was used. Using PACKMOL-Memgen, the bilayer system was prepared, solvated, and necessary counter ions were added. The minimum water distance from the membrane surface to the solvent box boundaries was increased to 35 Å to leave enough space between the substrate and the membrane surface and avoid interactions with periodic images during the extraction. Box dimensions in the x and y axes were set to 70 Å, resulting in systems comprised of ∼50,000 atoms.

### Simulated extraction of substrates from the membrane

MD simulations were performed using the GPU implementation of the AMBER 16 molecular simulation package [87, 88], employing the ff14SB force field for the protein [89], the Lipid17 force field for the lipids [90–92], and the TIP3P water model [93]. The SHAKE algorithm [94] was used to constrain bond lengths of hydrogen atoms to heavy atoms, enabling a time step of 2 fs. Long-range electrostatic interactions were considered using the Particle Mesh Ewald (PME) algorithm [95]. The system was energy-minimized by three mixed steepest descent/conjugate gradient calculations with a maximum of 20,000 steps each. First, the initial positions of the protein and membrane were restrained, followed by a calculation with restraints on the protein atoms only, and finalizing with a minimization without restraints. The minimized system was then gradually thermalized in two stages. Initially, the temperature was increased from 0 K to 100 K under NVT conditions, then from 100 K to 300 K under NPT conditions at 1 bar, using a Langevin thermostat [96]. The equilibration process continued for 5 ns, before starting with production simulations. As usual in membrane MD simulations, the NPT ensemble was used, allowing the membrane to accommodate along the trajectory [97]. For US simulations, the pressure was maintained using an anisotropic Berendsen barostat [98], while for the rest of the simulations a semi-isotropic Berendsen barostat [98] was used, coupling the membrane (x-y) plane with the constant-surface-tension dynamics. All analyses were performed by using CPPTRAJ [99]. Unless otherwise stated, molecular visualization was performed with Pymol [100] and VMD [101]. The Movie maker module within VMD was used to illustrate the acyl chain mobility and the access of substrates into PlaF.

To extract a substrate molecule from the membrane into one of the access tunnels, we selected the lipid that was closest to the entrance and pulled it from the membrane through the tunnel to the active site of PlaF, using constant velocity sMD simulations. Pulling simulations at low velocities are recommended for small polar molecules [102] and large lipids [103] to calculate free energy profiles. At the lowest pulling rates, lipids have time to adapt to energetically favorable conformations during the extraction process [103]. In a recent study investigating GPL binding to phospholipase A2 (PLA_2_), a constant pulling velocity of 5 Å ns^-1^ was used [57]. For the extraction of substrates, we considered all three possibilities by which a substrate may enter a tunnel: either the head group or one of the two tails. Depending on the type of head group (i.e., PG or PE), each substrate was pulled by its oxygen or nitrogen atoms at a constant velocity of 1 Å ns^-1^ using a force constant of 5 kcal mol^-1^ Å^-2^. When pulling at the tail, the terminal carbon atom of the respective acyl chain was used.

Each tunnel was divided into several segments connected through virtual points formed by the COM of amino acids lining the respective tunnel. The number of virtual points depends on the length and shape of the respective tunnel. The virtual points guided the extraction of substrates such that the substrates followed the path of the respective tunnel. In addition, to obtain a low energy pathway, an adaptive sMD protocol was implemented. For this, 50 replicas for each pulling simulation were carried out, and the work required was computed as a function of the reaction coordinate. The computed work was further related to free energy difference between two states of the pulling simulation applying Jarzynski’s relation (eq. 1) [30].

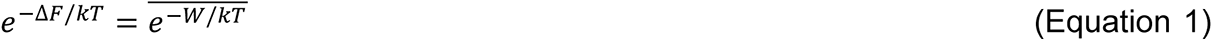

Here, Δ*F* is the free energy difference between two states, which is connected to work *W* done on the system [30]. *k* is the Boltzmann constant and *T* the temperature of the system. The replica closest to the Jarzynski’s average [30] was considered to describe the lowest-free-energy pathway and provided the starting point for the next pulling stage. Trajectories further away from that pathway were removed. This procedure results in faster convergence of PMF profiles, decreasing the overall computation needed [31].

For the systems without t-PlaF_A_, the substrates were extracted with the same pulling velocity and spring constant, as mentioned above. However, to avoid edge effects, a substrate in the middle of the membrane was located. For this extraction process, the reaction coordinate was the distance between the head atom of the pulled substrate and COM of phosphorous atoms of the lipids in the opposite leaflet. Furthermore, to determine the free energy minimum of the phospholipids in the membrane more accurately, the substrate was first pulled into the membrane (∼3 Å), before pulling it out of the membrane.

### Umbrella sampling and potential of mean force calculations

To understand the substrate access mechanism in PlaF and to identify preferential substrate access tunnels, PMFs were computed based on US [34], taking structures from the sMD simulations as starting points. As a reaction coordinate, the COM distance of the three oxygen atoms of the glycerol moiety in the substrate to the COM of residues S137 and H286 (only Cα atoms) of the active site was used. This reaction coordinate was also taken for over all other systems for it describes the essential aspects of the structural transformation during substrate access. Consecutive positions of the substrates from the membrane to the active site as determined by pulling simulations were considered reference points for US, with each position corresponding to one umbrella window. To achieve sufficient overlap between the umbrella windows, distances between reference points of ∼1 Å were used along the reaction coordinate. The length of individual tunnels and the size of acyl chains for respective substrates vary. Therefore, for sampling the access of different substrates, different numbers of windows were required for each tunnel. Selected positions of the lipid in the tunnel were restrained by harmonic potentials, using a force constant of 5 kcal mol^-1^ Å^-2^. To achieve sufficient convergence of the PMF profile, each window was sampled for 300 ns, of which the last 100 ns were used to calculate the PMF. Distance values were recorded every 2 ps and processed with WHAM [35, 36]. To estimate the PMF error, the data was separated into blocks according to the maximum calculated autocorrelation time of 20 ns. The correlation time was obtained for the complete trajectory, excluding the first 20 ns of sampling data for equilibration. The last 100 ns of sampling data was split into five blocks of 20 ns each, a PMF profile was calculated for each block with WHAM, and the error at each PMF point was calculated as the standard error of the mean.

Similarly, for systems without t-PlaF_A_, trajectories obtained by pulling simulations were used to set up US simulations. Umbrella windows were extracted at distances of 1 Å from the starting point of the pulling simulation until the substrate was not interacting with the membrane anymore. The selected positions of the lipid were restrained by harmonic potentials, using a force constant of 5 kcal mol^-1^ Å^-2^ and as the reaction coordinate the distance of the COM of the three oxygen atoms of the glycerol moiety of the substrate to the COM of phosphorous atoms of the lower membrane leaflet. Each window was simulated for 100 ns at constant pressure (1 bar) and temperature (300 K) conditions until convergence was achieved. The first 20 ns of simulation data was discarded. WHAM [35, 36] was used to calculate the PMF. The PMFs were evaluated for convergence by checking the change in the free energy profile with the increase in sampling time at every 10 ns. Furthermore, histograms of sampled configurations were visually inspected for sufficient overlap between the neighboring umbrella windows; otherwise, the iterative cycle in WHAM fails to converge and the free energy profiles have discontinuities.

### Absolute binding free energy from computed PMF

The absolute binding free energy of substrates to PlaF was determined from the computed PMF using an approach modified from Chen and Kuyucak [104]. The PMF was integrated along the reaction coordinate (eq. 2) to calculate an association (equilibrium) constant (*K*_eq_).

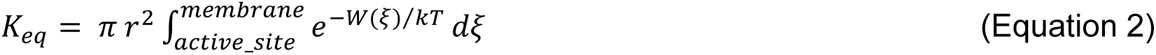

Here, *r* is the maximum bottleneck radius of the respective tunnel, which was determined by a CAVER analysis (Table 1), π*r*^2^ is the cross-sectional area of the tunnel, *W*(*ξ*) is the PMF at a specific value of the reaction coordinate, *k* is the Boltzmann constant, and *T* is the temperature at which the simulations were performed.

*K*_eq_ was then transformed to the mole fraction scale (*K*_x_), taking into account the number of lipids (*N*_L_) per surface area *A* (eq. 3).

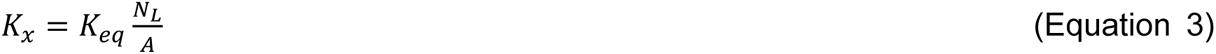

From *K*_x_, the difference in the free energy (eq. 4) between the bound and unbound state 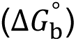 of a single substrate molecule was calculated.

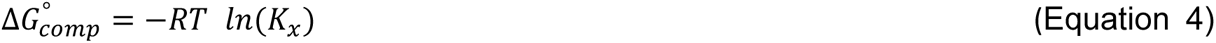

### Blocking access of the PlaF substrates

To corroborate predicted access tunnels for PlaF substrates, we intended to block these by small-to-tryptophan substitutions of tunnel-lining residues. To do so, we identified possible substitution sites from our previous CAVER analyses, taking into account the tunnels’ bottleneck radii and lengths. For these analyses, the same trajectory used to search for tunnels in t-PlaF_A_ was considered. Finally, 4-5 amino acids within each tunnel were selected for substitutions.

In the first step, all the amino acids except glycines and prolines within 3 Å of individual tunnels and oriented towards a tunnel were considered. In turn, residues with an outward orientation were disregarded as a substitution there will likely not block the tunnel. Furthermore, as the TM and JM helix was found to be important for both the dimerization and the activity of PlaF [19], residues of these helices were excluded. Finally, the catalytic residues S137, D258, and H286 and other residues of the active site were disregarded to avoid affecting the activity of PlaF.

The selected residues of each tunnel were substituted to tryptophan using FoldX [105], and the stability of the Plaf variants was evaluated in terms of the change in free energy (ΔΔ*G*) with respect to the wild type [106]. Single amino acid substitutions were performed 10 times for each proposed residue of each tunnel, and the results were averaged. If the average ΔΔ*G* > 3 kcal mol^-1^, the substitution is considered destabilizing [107] and was not further pursued. To check if the proposed substitutions will block the tunnel, the bottleneck radius of the variant tunnels was recalculated using CAVER. As done earlier, the probe radius was set to 2 Å. If no tunnel was identified with this criterium, the probe radius for tunnel search was reduced until the tunnels started to appear again.

### Biological evaluation of PlaF activity from mutations

#### a) Site-directed mutagenesis, protein expression, and purification

The plasmids for expression of PlaF variants with substitutions in the tunnels (Table S5) were created by PCR, using Phusion DNA polymerase (Thermo Fischer Scientific) in whole plasmid amplification, with mutagenic oligonucleotides (Table S7) designed for the SLIC method [108], and p-*plaF* plasmid [19] as a template. The presence of desired nucleotide substitutions was confirmed by DNA sequencing (MWG Biotech, Ebersberg, Germany). PlaF was purified from *P. aeruginosa* p-*plaF* membranes and solubilized with DDM, as described previously [109]. Proteins were analyzed by polyacrylamide gel electrophoresis under denaturation conditions (SDS-PAGE) on 14% (w/v) gels, as described by Laemmli [110]. The protein concentration was determined by measuring the *A*_280nm_ using a NanoDrop 2000C spectrophotometer (Thermo Fisher Scientific Inc., Waltham, Massachusetts, USA). The extinction coefficients for PlaF and the variants were calculated with the ProtParam tool (Navia-Paldanius *et al.*, 2012), considering the amino acid exchange and a His_6_-tag.

#### b) Enzyme activity assays and kinetic studies

The esterase activities of PlaF and variants were determined with *p*-NPB as substrate as described previously [111], using a 96-well microplate and starting the reaction by adding 100 µl of PlaF sample (16 nM) to the 100 µl of *p*-NPB solution (2 mM). Kinetic parameters (*K*_m_ and *k*_cat_) for hydrolysis of *p*-NPB were determined using 8 nM enzyme as described previously [109]. Kinetic parameters were determined by non-linear regression analysis of data fitted to the Michaelis-Menten equation with PrismLab. Enzyme activities with DLPG and LGPLs were determined according to the established protocol in ref. [112]. For enzymatic reactions, 25 µL of PlaF or the variant (16 nM) and 25 µL of DLPG solution were used. The amount of released FAs after 24 h of reaction were calculated from the calibration curve using oleic acid at concentrations ranging from 0.1 to 1.0 mM.

#### c) Thermal stability

PlaF and variants (28.1 µM, 10 µL) loaded into the measuring capillaries (Prometheus NT.Plex nanoDSF Grade Standard Capillary Chips) were heated from 20 to 90 °C (1 °C min^-1^ heating rate), and the intrinsic protein fluorescence was recorded at 330 nm and 350 nm using the Prometheus NT.Plex nanoDSF device (Nano Temper, Munich, Germany) [113]. The melting points were calculated from the first derivative of the ratio of *F*_350nm_ and *F*_330nm_ using the PR.ThermControl software (Nano Temper, Munich, Germany) [113].

### Egress of PlaF products

To determine the egress pathways of PlaF products, a system with 2LMG substrate was considered. The final snapshot at 300 ns of the US simulations of the window with the substrate close to the active site was considered as the starting structure for unbiased MD simulations. 2LMG was cleaved into the products: MYR and PGR, without altering the orientation of each product within the tunnels. Atomic partial charges for the products were derived according to the restraint electrostatic potential fit (RESP) procedure [114], as implemented in Antechamber [115]. Geometry optimizations and subsequent single-point calculations were performed with Gaussian [116] at the Hartree-Fock (HF) level with the 6-31G* basis set. Force field parameters for the products were taken from the general amber force field for organic molecules (GAFF, version 2) [117]. The prepared system was then minimized, thermalized, and equilibrated using the protocol described above for MD simulations. 12 replicas of production MD simulations of 3 µs length each under NPT conditions were performed. The distance of the 2LMG products to the entrance of each tunnel was computed for each replica.

## Acknowledgments

This study was funded by the Deutsche Forschungsgemeinschaft (DFG, German Research Foundation) project no. 267205415 / CRC 1208 grant to FK and KEJ (TP 02) as well as HG (TP 03). We are grateful for computational support by the “Zentrum für Informations und Medientechnologie” at the Heinrich-Heine-Universität Düsseldorf and the computing time provided by the John von Neumann Institute for Computing (NIC) to HG on the supercomputer JUWELS at Jülich Supercomputing Centre (JSC) (user IDs: HDD18; plaf).

## Author contributions

H.G. conceptualization, supervision, analysis, visualization, writing; F.K. conceptualization, supervision, analysis, visualization, writing; K.-E.J. supervision, writing; S.A. investigation, analysis, visualization, writing; C.H.S. investigation, analysis, visualization, writing; S.N.S.V. analysis, visualization, writing.

## Additional information

Supplementary information is available for this paper.

