## Supplemental Information for "Substrate access mechanism in a novel membrane-bound phospholipase A of *Pseudomonas aeruginosa* concordant with specificity and regioselectivity"

### Table of Contents

1. Supplementary results ........................................................................................S3
2. Supplementary tables ........................................................................................S5
3. Supplementary figures ........................................................................................S10
4. Supplementary movies ........................................................................................S27
5. Supplementary references ........................................................................................S28

### 1. Supplementary results

##### Energetics of GPL and LGPL extraction from the membrane

As reference points, we computed the energetics of extracting a single PlaF substrate from the membrane to the solvent. We chose 1,2-dilauroyl-*sn*-glycero-3-phosphoglycerol (DLPG) and 1-myristoyl-2-hydroxy-*sn*-glycero-3-phosphoglycerol (2LMG), the two glycerophospholipid (GPL) and lysoglycerophospholipid (LGPL) substrates for which PlaF shows the highest activity [1], and also see main text, Figure 3. Applying Steered Molecular Dynamics (sMD) simulations, the substrates were pulled out of the membrane with their head group first, applying a constant velocity of 1 Å $ns^{-1}$, until the hydrophobic tails do not interact anymore with the membrane surface. Once in the solvent, the hydrophobic chains curl in order to minimize their exposure to water.

The differences in the potentials of mean force (PMF) between the states in the solvent and in the membrane are ~13 ± 0.1 kcal $mol^{-1}$ for DLPG and ~8 ± 0.3 kcal $mol^{-1}$ for 2LMG (Figure S2A). Converged PMFs of extraction of the substrates were obtained after ~40 ns of umbrella sampling (US) simulation time per window (Figure S2B), which also resulted in sufficient overlap between the reaction coordinate distributions of neighboring windows (Figure S2C).

Unfortunately, the PMF cannot be measured experimentally. However, the free energy difference between the two states of the lipid in the solvent and in the membrane can be related to the experimentally measured critical micelle concentration (CMC) [2]. The CMC can be related to an excess chemical potential (eq. S1).

$$\mu - \mu_0 = R * T * \ln\left(\frac{CMC}{55.5\ M}\right) \quad \text{Equation S1}$$

$\mu - \mu_0$ is the excess chemical potential, *R* is the gas constant, *T* is the temperature, and CMC has been converted to mole fraction units. The CMC values were obtained from previous studies [3, 4]. The determined excess chemical potential for DLPG and 2LMG is 11.69 kcal $mol^{-1}$ and 7.60 kcal $mol^{-1}$, respectively. These values are within chemical accuracy [5, 6] to those obtained from the PMF.

##### Energetics of substrate access into T2 in dimeric PlaF

In dimeric PlaF (di-PlaF), the orientation of the tunnels with respect to the membrane changes (Figure S18A), and the tunnel entrances are higher above the membrane interface. In this configuration, T2 is closest of all tunnels to the membrane interface with a distance of 7.4 ± 1.5 Å (Figure S18A). Therefore, we computed the PMF for substrate access across T2 in di-PlaF. We follow the same steps considered for the substrate access in tilted monomeric PlaF (t-PlaF$_A$). The tail 1 access of DLPG in di-PlaF revealed a free energy barrier of 13 kcal mol$^{-1}$ (Figure S18B), compared to no energy barrier in t-PlaF$_A$ (see the main text, Figure 4B). The PMF was found converged after 300 ns yielding a maximal difference of ~0.5 kcal mol$^{-1}$ as to a PMF computed from 280 ns per window (Figure S18C), and neighboring umbrella windows have sufficient median overlap of 4.2% (Figure S18C). These results indicate that substrate access via tail 1 across T2 is disfavorable in di-PlaF compared to that in t-PlaF, and may explain why PlaF is inactive in the dimeric configuration [1].

**Computational costs to determine PMFs related to substrate access**

Considering the different substrates and their modes of access across the tunnels T1 and T2, there are 11 systems for which we calculated the PMF from US simulations. Depending on the properties of the substrates, their position in the membrane, their access mode, and the tunnel, each system required a different number of umbrella windows, resulting in different computational costs. In total, all computations add up to ~104 µs of sampling simulations for the substrates investigated in this study (Table S8).

#### 2. Supplementary tables

**Table S1:** Characteristics of tunnels identified in $PlaF_A$, $PlaF_B$, and di-PlaF using CAVER.

| Tunnel | Occurrence[a,b] | | | | Average length[c] | | | |
|---|---|---|---|---|---|---|---|---|
| | Monomer | | Dimer | | Monomer | | Dimer | |
| | $PlaF_A$ | $PlaF_B$ | di-$PlaF_A$ | di-$PlaF_B$ | $PlaF_A$ | $PlaF_B$ | di-$PlaF_A$ | di-$PlaF_B$ |
| T1 | 9.12 | 6.07 | 3.30 | 5.55 | 25.72 | 23.90 | 41.07 | 23.81 |
| T2 | 18.35 | 5.07 | 18.45 | 5.02 | 23.97 | 22.35 | 23.89 | 22.20 |
| T3 | 16.80 | 6.25 | 21.30 | 5.80 | 15.84 | 16.16 | 15.52 | 15.78 |

[a] Snapshots in which the tunnel is identified with respect to the total number of snapshots, in %.
[b] Data calculated with a probe radius of 2.0 Å.
[c] In Å.

**Table S2:** Pulling points across the tunnels for sMD simulations.

| T1 pulling points[a,b] | Amino acid residues | T2 pulling points[a,c] | Amino acid residues |
|---|---|---|---|
| A1 | V30, P205, L206 | B1 | A24, S102 |
| A2 | E34, F192 | B2 | L27, N225 |
| A3 | G72, L214, V287 | B3 | D76, F192, N225 |
| A4 | F71, D161, F192 | B4 | A73, V199, A221 |
| A5 | M138, L184, H286 | B5 | F71, D161, F192 |
| A6 | M138, F174 | B6 | M138, L184, H286 |
| A7 | K170, Q234, Y236 | B7 | M138, F174 |
| A8[d] | S137 | B8 | K170, Q234, Y236 |
| | | B9[d] | S137 |

[a] Pulling points are COM of corresponding amino acid residues.
[b] For T1, A1-A4 are components of T1, and A5-A8 are components of T3.
[c] For T2, B1-B4 are components of T2, B5 is a component of T1, and B6-B8 are components of T3.
[d] For S137 of A8/B9, the OH group of the nucleophile was considered as a pulling point.

**Table S3:** Overview of sMD simulations for the substrate access through T1 and T2 in t-PlaF$_A$.

| Tunnel | Substrate | Mode of access | Per access simulation time[a] | Number of replicas | Total simulation time[b] | Per substrate simulation time[b] |
|---|---|---|---|---|---|---|
| T1 | DLPG | head | ~46 | 50 | ~2.30 | ~8.80 |
| | | tail 1 | ~62 | 50 | ~3.10 | |
| | | tail 2 | ~67 | 50 | ~3.35 | |
| | DLPE | tail 1 | ~60 | 50 | ~3.00 | ~6.00 |
| | | tail 2 | ~60 | 50 | ~3.00 | |
| | DSPG | tail 1 | ~66 | 50 | ~3.30 | ~6.75 |
| | | tail 2 | ~69 | 50 | ~3.45 | |
| | 2LMG | head | ~51 | 50 | ~2.55 | ~5.00 |
| | | tail 1 | ~49 | 50 | ~2.45 | |
| T2 | DLPG | head | ~57 | 50 | ~2.85 | ~10.25 |
| | | tail 1 | ~70 | 50 | ~3.50 | |
| | | tail 2 | ~78 | 50 | ~3.90 | |
| | DLPE | tail 1 | ~54 | 50 | ~2.70 | ~5.75 |
| | | tail 2 | ~61 | 50 | ~3.05 | |
| | DSPG | tail 1 | ~63 | 50 | ~3.15 | ~6.15 |
| | | tail 2 | ~60 | 50 | ~3.00 | |
| | 2LMG | head | ~52 | 50 | ~2.60 | ~5.30 |
| | | tail 1 | ~54 | 50 | ~2.70 | |

[a] In ns.
[b] In µs.

**Table S4:** Overview of computed absolute binding free energy of DLPG to t-PlaF$_A$ from PMF.

| **System** | T1HG | T1T1 | T1T2 | T2HG | T2T1 | T2T2 |
|---|---|---|---|---|---|---|
| $\Delta G^{\circ}_{comp}$ [a,b] | 1.81 ± 0.25 | 1.31 ± 0.16 | -4.01 ± 0.56 | 1.00 ± 0.48 | -2.89 ± 1.46 | 0.96 ± 0.27 |

[a] In kcal mol$^{-1}$.
[b] Error estimation: For each system, the last 100 ns of sampling data was split into five independent blocks of 20 ns each. The PMF profiles obtained were used to determine the absolute binding free energy for each block, and the standard error of the mean was calculated.

**Table S5:** Structural stability of proposed tunnel variants of PlaF determined using FoldX and corresponding influence on tunnel characteristics calculated with CAVER.

| **Tunnel** | **PlaF variant** | **ΔΔG[a,b]** | **Average bottleneck radius[c,d]** | **Average length[d]** |
|---|---|---|---|---|
| T1 | N77W | -0.48 | 2.21 | 27.40 |
| | R80W | 0.68 | 2.42 | 26.08 |
| | L214W | 1.13 | 2.15 | 26.84 |
| | V290W | -0.65 | 1.83 | 26.06 |
| T2 | D74W | -0.69 | 1.63 | 30.85 |
| | R217W | -0.10 | 1.87 | 24.30 |
| | A218W | 0.40 | 1.86 | 23.67 |
| | A221W | -0.11 | 1.82 | 28.26 |
| | N225W | -1.20 | 1.80 | 29.77 |
| T3 | M166W | 0.15 | 2.11 | 14.72 |
| | L177W | 0.48 | 2.24 | 14.25 |
| | F229W | -0.28 | 2.23 | 14.18 |
| | R233W | -0.50 | 2.16 | 18.54 |
| | Y236W | -0.18 | 2.10 | 13.82 |

[a] $\Delta\Delta G = \Delta G_{\text{variant}} - \Delta G_{\text{wild type}}$.
[b] In kcal mol$^{-1}$.
[c] Data calculated with a probe radius of 1.2 Å.
[d] In Å.

**Table S6:** Statistical test[a] to determine the tendency of MYR reaching the entrance of tunnels T1-T3 in 12 replicas.

| **Tunnels** | **T3** | **T2** |
|---|---|---|
| **T1** | 2.5981 ($p$ = 0.0047) | 3.1436 ($p$ = 0.0008) |
| **T2** | -1.0215 ($p$ = 0.1539) | |

[a] The z-score for two population proportions related to two tunnels was calculated [7-9]. A cutoff of 5 Å was chosen to identify the tunnels where MYR reaches the entrance during 12 independent replicas of 3 μs long unbiased MD simulations. In 7 replicas MYR reaches T1, in 1 replica T3, but it does not reach the T2 entrance (Figure S17). The tendency of MYR reaching the entrance of T1 is significantly higher (at $p < 0.05$, considering a one-tailed z-score test) than reaching the entrance of T2 or T3.

**Table S7:** List of the oligonucleotides used for generating expression plasmids.

| Oligonucleotide | DNA sequence (5'→3')[a] | Plasmid |
|---|---|---|
| PlaF$_{D74W}$-up | CTTCGGCGCC**TGG**AAGGACAACTGGCTGCGCTTCGC | p-*plaF*$_{D74W}$ |
| PlaF$_{D74W}$-down | GTTGTCCTT**CCA**GGCGCCGAAGCCGTGGATCAGCAAC | |
| PlaF$_{N77W}$-up | GACAAGGAC**TGG**TGGCTGCGCTTCGCCCGGCCGCTG | p-*plaF*$_{N77W}$ |
| PlaF$_{N77W}$-down | GCGCAGCCA**CCA**GTCCTTGTCGGCGCCGAAGCCGTG | |
| PlaF$_{R80W}$-up | AACTGGCTG**TGG**TTCGCCCGGCCGCTGACCGAGCG | p-*plaF*$_{R80W}$ |
| PlaF$_{R80W}$-down | CCGGGCGAA**CCA**CAGCCAGTTGTCCTTGTCGGCGC | |
| PlaF$_{M166W}$-up | GCCGGGGTG**TGG**CCGGCGCGCAAGAGCGAACTGTTC | p-*plaF*$_{M166W}$ |
| PlaF$_{M166W}$-down | GCGCGCCGG**CCA**CACCCCGGCGTTGTCGATCAGCG | |
| PlaF$_{L177W}$-up | GTTCGAGGAC**TGG**GAGCGCGGCGAGAATCCCCTGGTG | p-*plaF*$_{L177W}$ |
| PlaF$_{L177W}$-down | GCCGCGCTC**CCA**GTCCTCGAACAGTTCGCTCTTGC | |
| PlaF$_{L214W}$-up | CAAGCGCTAC**TGG**GGCGAGCGCGCGGTAGCCGCGTC | p-*plaF*$_{L214W}$ |
| PlaF$_{L214W}$-down | GCGCTCGCC**CCA**GTAGCGCTTGAGCGGCGCCGGCAG | |
| PlaF$_{R217W}$-up | CTCGGCGAG**TGG**GCGGTAGCCGCGTCGGCGTTCAAC | p-*plaF*$_{R217W}$ |
| PlaF$_{R217W}$-down | GGCTACCGC**CCA**CTCGCCGAGGTAGCGCTTGAGCG | |
| PlaF$_{A218W}$-up | GGCGAGCGC**TGG**GTAGCCGCGTCGGCGTTCAACGC | p-*plaF*$_{A218W}$ |
| PlaF$_{A218W}$-down | CGCGGCTAC**CCA**GCGCTCGCCGAGGTAGCGCTTGAG | |
| PlaF$_{A221W}$-up | GCGGTAGCC**TGG**TCGGCGTTCAACGCGCAGATATTC | p-*plaF*$_{A221W}$ |
| PlaF$_{A221W}$-down | GAACGCCGA**CCA**GGCTACCGCGCGCTCGCCGAGGTAG | |
| PlaF$_{N225W}$-up | GTCGGCGTTC**TGG**GCGCAGATATTCGAACAACTGCG | p-*plaF*$_{N225W}$ |
| PlaF$_{N225W}$-down | GAATATCTGCGC**CCA**GAACGCCGACGCGGCTACGCGC | |
| PlaF$_{R233W}$-up | GAACAACTG**TGG**CAGCGCTACATCCCGCTGGAGCC | p-*plaF*$_{R233W}$ |
| PlaF$_{R233W}$-down | GTAGCGCTG**CCA**CAGTTGTTCGAATATCTGCGCGTTG | |
| PlaF$_{F229W}$-up | GCGCAGATA**TGG**GAACAACTGCGCCAGCGCTACATC | p-*plaF*$_{F229W}$ |
| PlaF$_{F229W}$-down | CAGTTGTTC**CCA**TATCTGCGCGTTGAACGCCGACG | |
| PlaF$_{Y236W}$-up | CGCCAGCGC**TGG**ATCCCGCTGGAGCCGGAACTGCC | p-*plaF*$_{Y236W}$ |
| PlaF$_{Y236W}$-down | CAGCGGGAT**CCA**GCGCTGGCGCAGTTGTTCGAATATC | |
| PlaF$_{V290W}$-up | GTGCCGATG**TGG**GAACGCCCGGAGGAAACCGCGCAG | p-*plaF*$_{V290W}$ |
| PlaF$_{V290W}$-down | CGGGCGTTC**CCA**CATCGGCACGTGTCCGCAGTTTTC | |

[a] Mutated sequence is indicated in bold.

**Table S8:** Setup of umbrella sampling simulations for the substrate access through T1 and T2.

| Substrate | Tunnel | Mode of access | No. of windows | Sampling length[a] | Total sampling length[a] |
|---|---|---|---|---|---|
| DLPG[b] | 1 | head | 28 | 0.3 | 8.4 |
| DLPG[b] | 1 | tail 1 | 32 | 0.3 | 9.6 |
| DLPG[b] | 1 | tail 2 | 35 | 0.3 | 10.5 |
| DLPG[b] | 2 | head | 34 | 0.3 | 10.2 |
| DLPG[b] | 2 | tail 1 | 37 | 0.3 | 11.1 |
| DLPG[b] | 2 | tail 2 | 34 | 0.3 | 10.2 |
| DSPG[b] | 2 | tail 1 | 32 | 0.3 | 9.6 |
| DLPE[b] | 2 | tail 1 | 26 | 0.3 | 7.8 |
| 2LMG[b] | 2 | head | 28 | 0.3 | 8.4 |
| 2LMG[b] | 2 | tail 1 | 32 | 0.3 | 9.6 |
| DLPG[c] | 2 | tail 1 | 28 | 0.3 | 8.4 |

[a] In µs.
[b] Considering t-PlaF$_A$.
[c] Considering di-PlaF.

#### 3. Supplementary figures

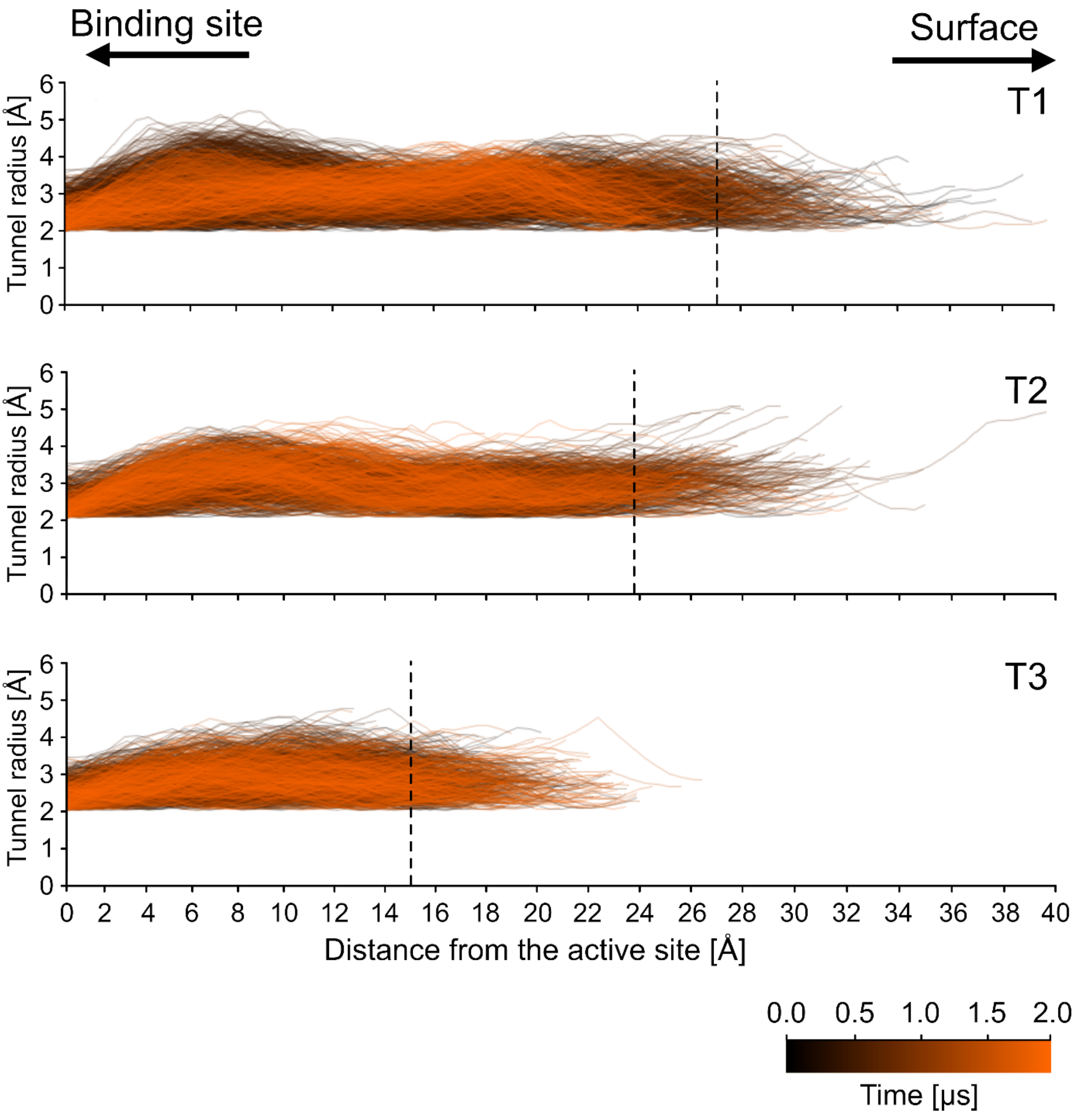


**Figure S1: Properties of tunnels identified in t-PlaF$_A$ ensembles.** Profiles of selected tunnel clusters T1-T3 were evaluated as to radius and distance from the active site during MD simulations of 2 µs length (see color scale). Each line represents the tunnel profile of a single snapshot. Black dashed lines mark the average length of tunnels in their respective cluster. T1 is the longest and T3 the shortest of the three tunnels.

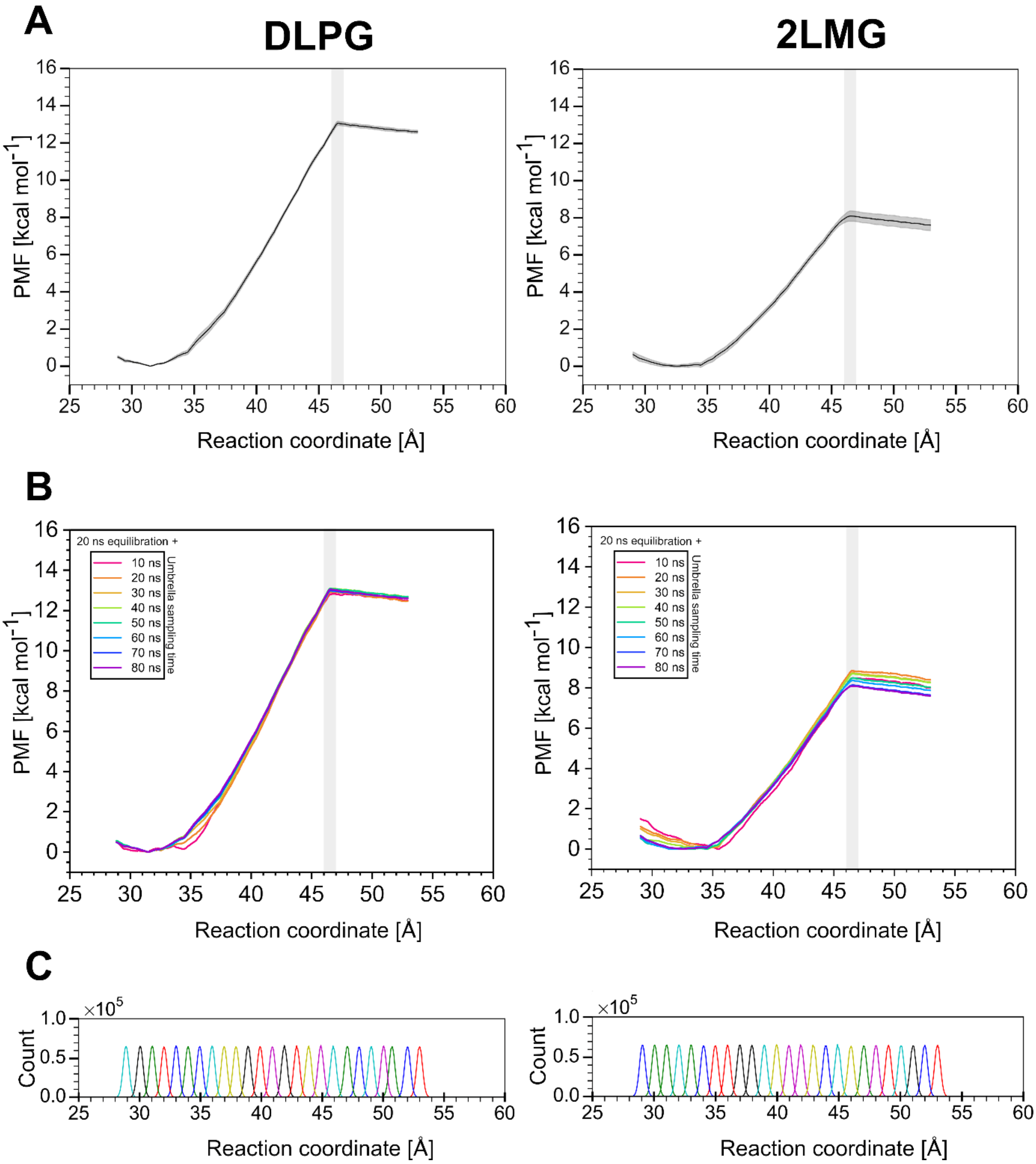


**Figure S2: Extraction of PlaF substrates from the membrane.** A) PMF profiles of selected substrates, DLPG (left) and 2LMG (right). B) Convergence plot indicates sufficient sampling time for the two substrates, DLPG (left) and 2LMG (right); convergence of profiles starts around 40 ns; at 100 ns, the PMF profiles are converged for both the substrates. C) Histograms indicate sufficient overlap among the umbrella windows of DLPG (left) and 2LMG (right) using a force constant of 5 kcal $mol^{-1}$ $Å^{-2}$; the median overlap is 4.13% and 4.10% for DLPG and 2LMG, respectively. The grey box in A and B indicates the section of the reaction coordinate where the substrate loses interaction with the membrane surface.

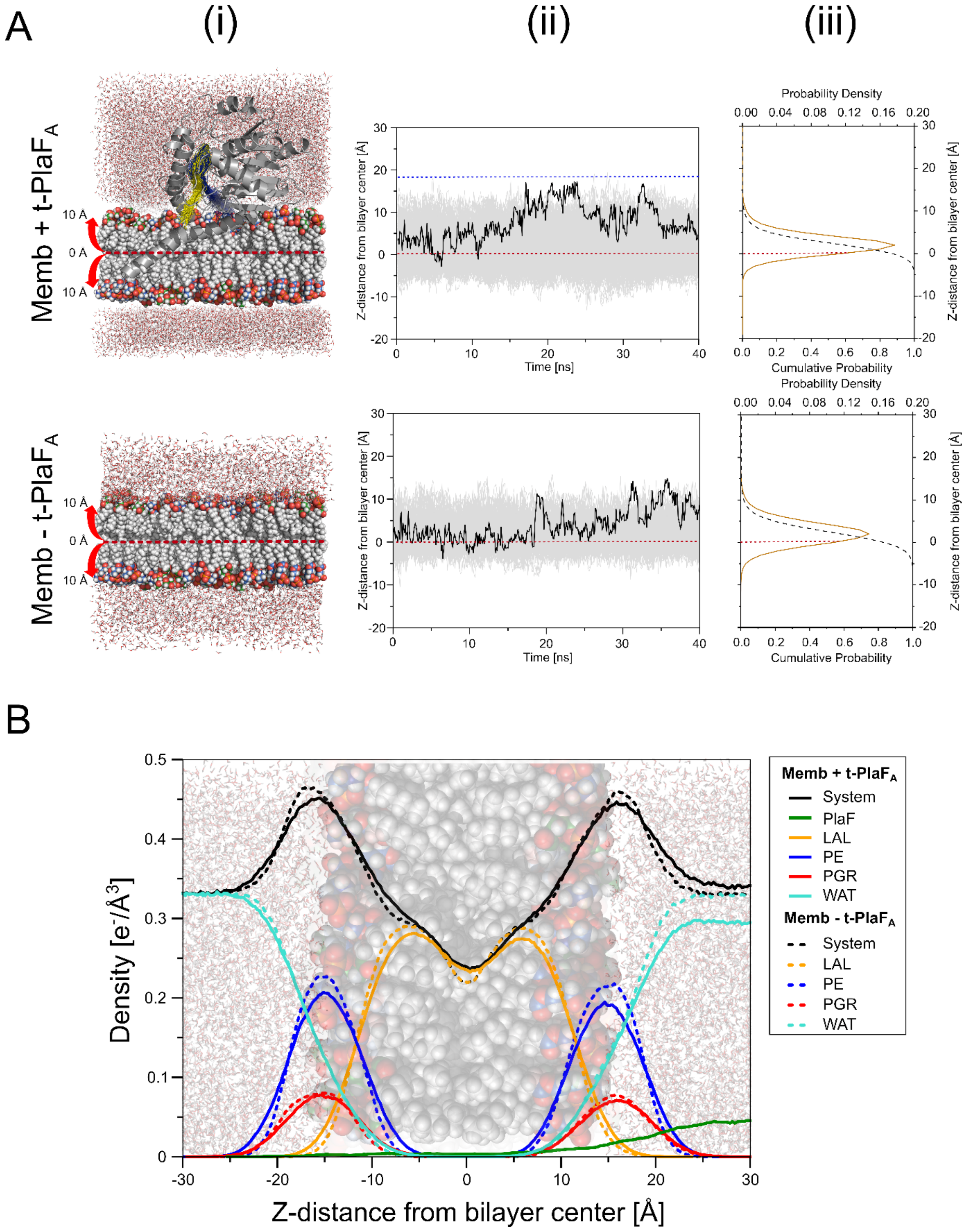


**Figure S3: Distribution of acyl chain termini from GPLs and corresponding density profiles.** A) To investigate the distribution of acyl chain termini during unbiased MD simulations, (i) a system with t-PlaF$_A$ (upper) and without t-PlaF$_A$ (lower) was considered. In t-PlaF$_A$, the tunnels T1 (blue) and T2 (yellow) are immersed into the head group region of the upper leaflet. (ii) With the bilayer center positioned at ≈ 0 Å (broken red line), the z-coordinate distance was measured for each acyl chain terminus during MD simulations of 40 ns. Tails from both upper and lower leaflets were considered. For both systems, the acyl chain termini can reach the membrane interface, located around 10-15 Å from the bilayer center (please see section B for the electron density profile). For the system with t-PlaF$_A$, the tail termini move as high as 20 Å, close to the entrance of access tunnels (broken blue lines) that are at z = 19.07 ± 1.42 Å for T1 and z = 17.94 ± 1.07 Å for T2. For the system without t-PlaF$_A$, the termini reach 16 Å of the z-coordinate. The acyl chain termini cannot only move up to the membrane surface, but also beyond the bilayer center along the negative z-coordinate. The black curve represents an example, where the acyl terminus of a selected lipid reaches the membrane interface (please see the movies S1 and S2 corresponding to this event for the systems with and without t-PlaF$_A$, respectively). iii) The probability density plot (brown curve) shows that the distribution of acyl chains shifts towards the positive z-coordinate, indicating that tails of GPLs can reach the membrane interface. The cumulative probability (broken black curve) of finding an acyl chain terminus at z > 10 Å is 1.5 % and 1.0 % for the systems with and without t-PlaF$_A$, respectively. B) The electron density profile was measured and compared for the two systems. Differences in the profiles are due to GPLs and water replaced by t-PlaF$_A$.

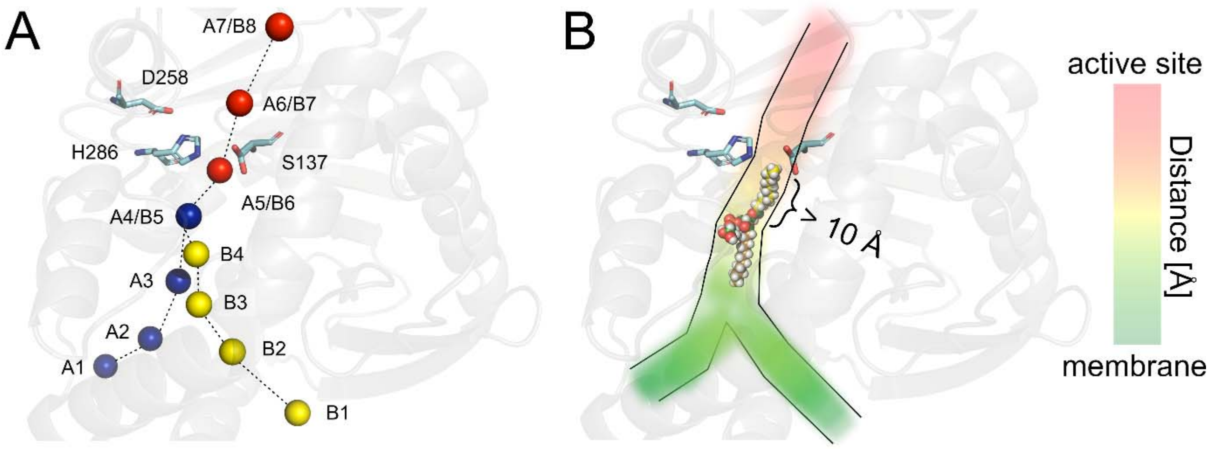


**Figure S4: Substrate access pathway.** A) Pulling points for substrate access in PlaF: using sMD simulations, substrates are first pulled out of the membrane to A1 (for T1, blue spheres) or B1 (for T2, yellow spheres). Red spheres correspond to pulling points lining T3. Substrate pulling through T1 involves points A1 to A7, while pulling through T2 involves points B1 to B8. T2 merges into T1 after A3; both follow a common path towards T3 across A4/B5. Catalytic residues are represented as cyan sticks. B) Requirement of T3 for substrate access: when pulled with terminal atoms, the *sn*-1 site of the substrate remains several Angstroms away from the catalytic S137 and, hence, needs to be further pulled into T3. Since the tunnels are almost straight, the reaction coordinate monotonically decreases as the substrate approaches the active site from the membrane.

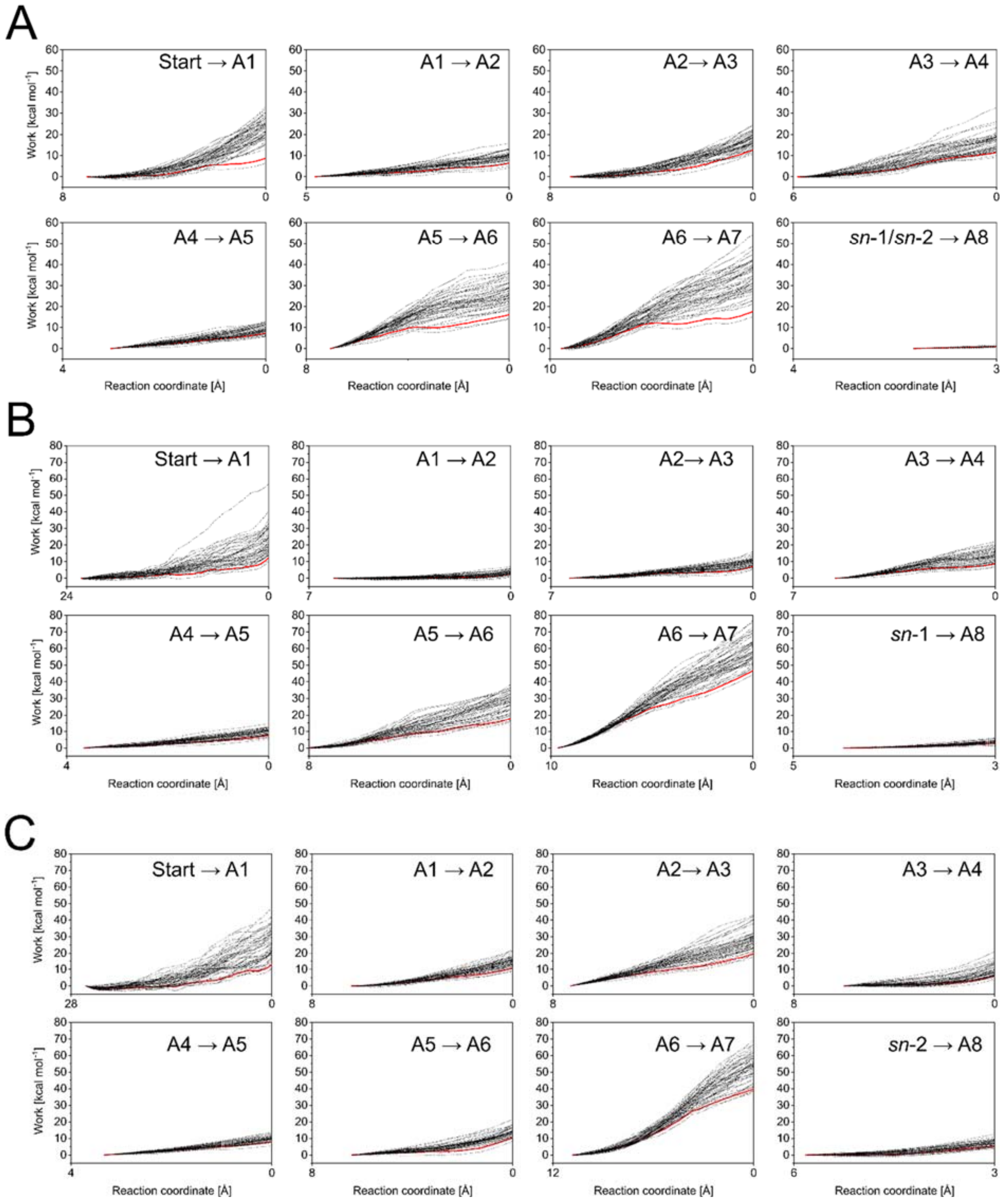


**Figure S5:** Work distributions (black lines) obtained from 50 replicas of sMD simulations to pull DLPG across T1 via (A) head, (B) tail 1, (C) tail 2. For each mode of access, DLPG is first pulled out of the membrane to point A1. A replica closest to Jarzynski's average (red line) was considered as the starting point for the next pulling, A1 → A2. This pulling continues until A7, after which the *sn*-1/*sn*-2 of DLPG is further pulled to the nucleophilic OH group of the catalytic S137. The reaction coordinate denotes the distance to the target point.

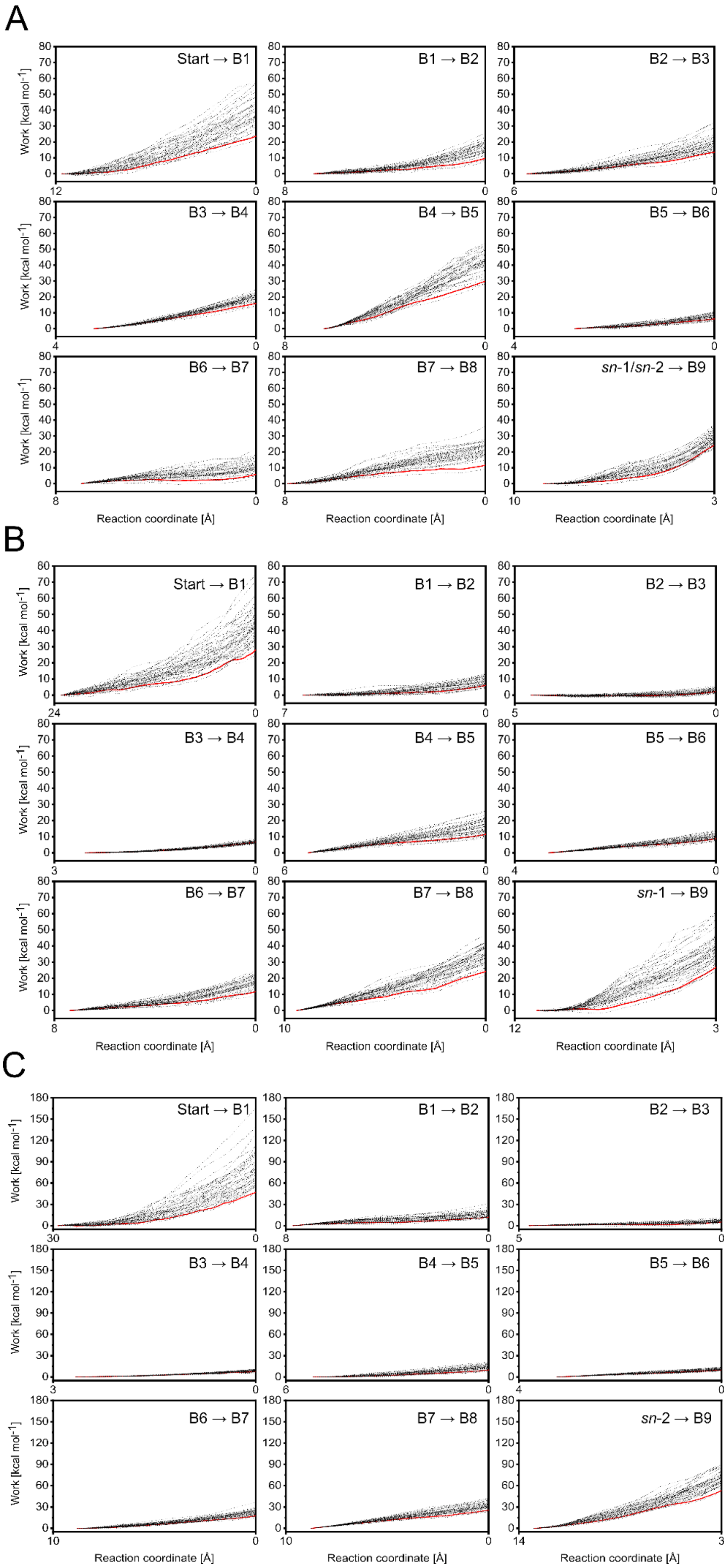


**Figure S6:** Work distributions (black lines) obtained from 50 replicas of sMD simulations to pull DLPG across T2 via (A) head, (B) tail 1, (C) tail 2. For each mode of access, DLPG is first pulled out of the membrane to point B1. A replica closest to Jarzynski's average (red line) was considered as the starting point for the next pulling, B1 → B2. This pulling continues until the pulling point B8, after which the *sn*-1/*sn*-2 of DLPG is further pulled to the nucleophilic OH group of the catalytic S137. The reaction coordinate denotes the distance to the target point.

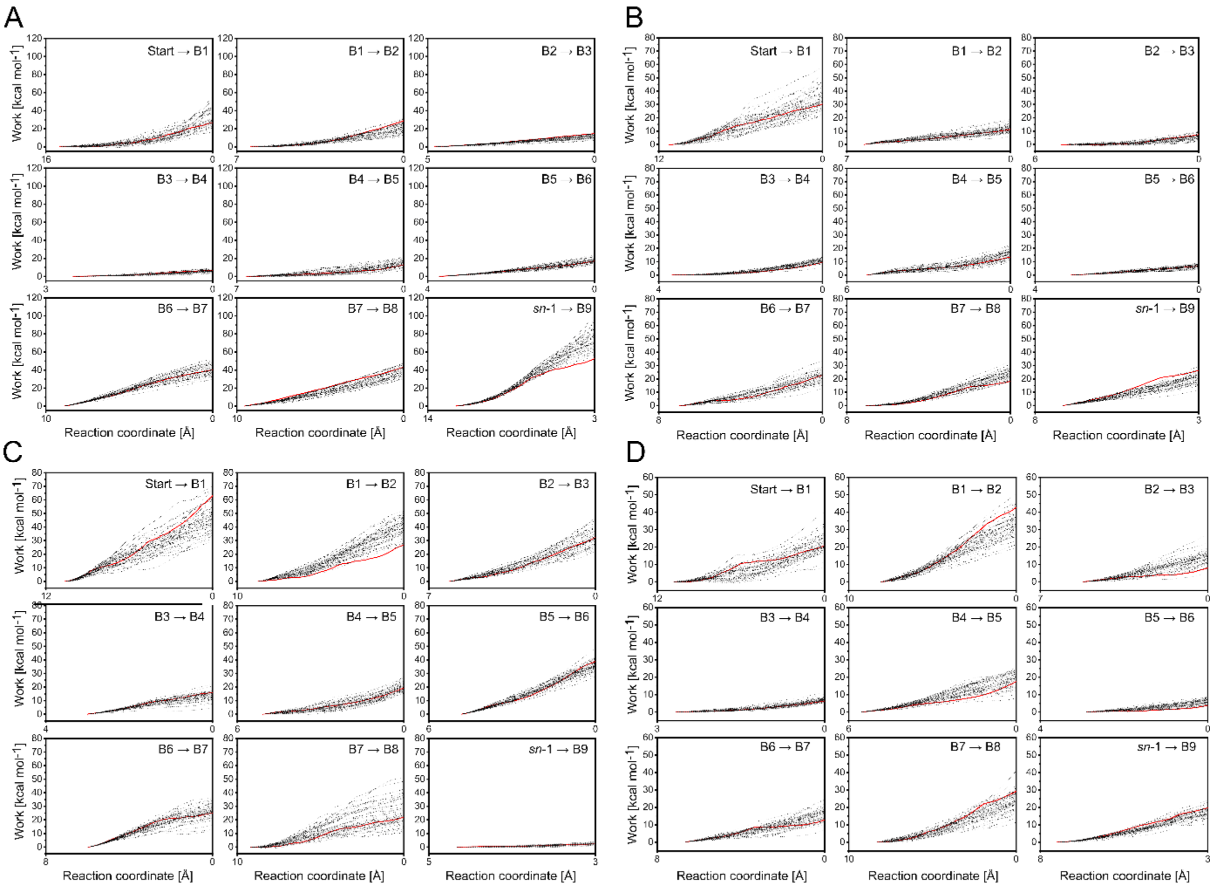


**Figure S7:** Work distributions (black lines) obtained from 50 replicas of sMD simulations to pull (A) DSPG via tail 1, (B) DLPE via tail 1, (C) 2LMG via head (D) 2LMG via tail 1 across T2. Each substrate was first pulled out of the membrane to point B1. A replica closest to Jarzynski's average (red line) was considered as the starting point for the next pulling, B1 → B2. This pulling continues until the pulling point B8, after which the *sn*-1 of respective substrate is further pulled to the nucleophilic OH group of the catalytic S137. The reaction coordinate denotes the distance to the target point.

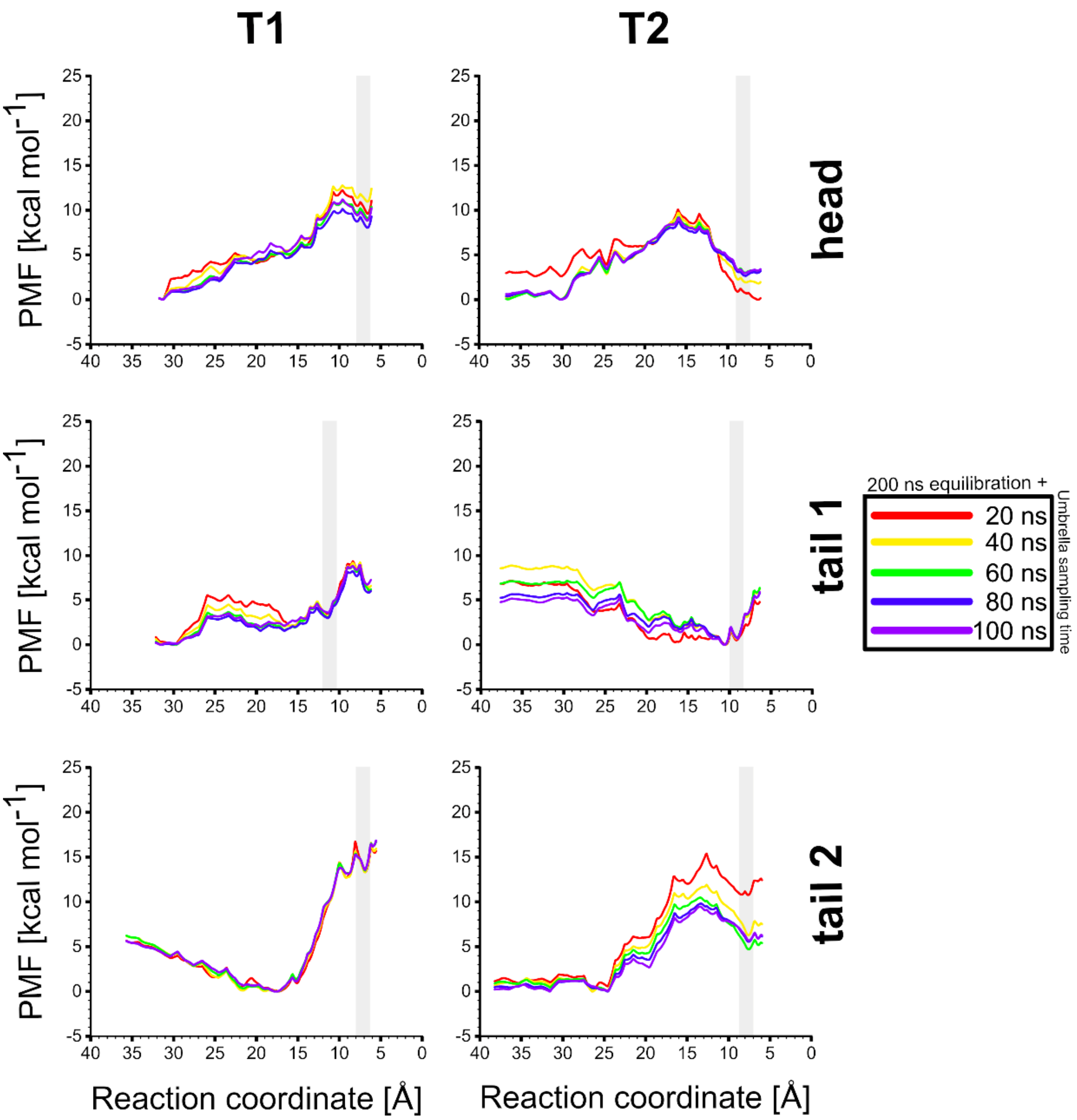


**Figure S8: Convergence of PMFs for substrate access of DLPG.** PMFs were computed every 20 ns for the range of 220-300 ns (see legend) of umbrella sampling simulations per window for T1 (left) and T2 (right). The first 200 ns of the sampling simulations were considered for equilibration and removed for every system. The grey box indicates the location of the active site. Overall, 300 ns of umbrella sampling per window are sufficient to achieve converged PMFs.

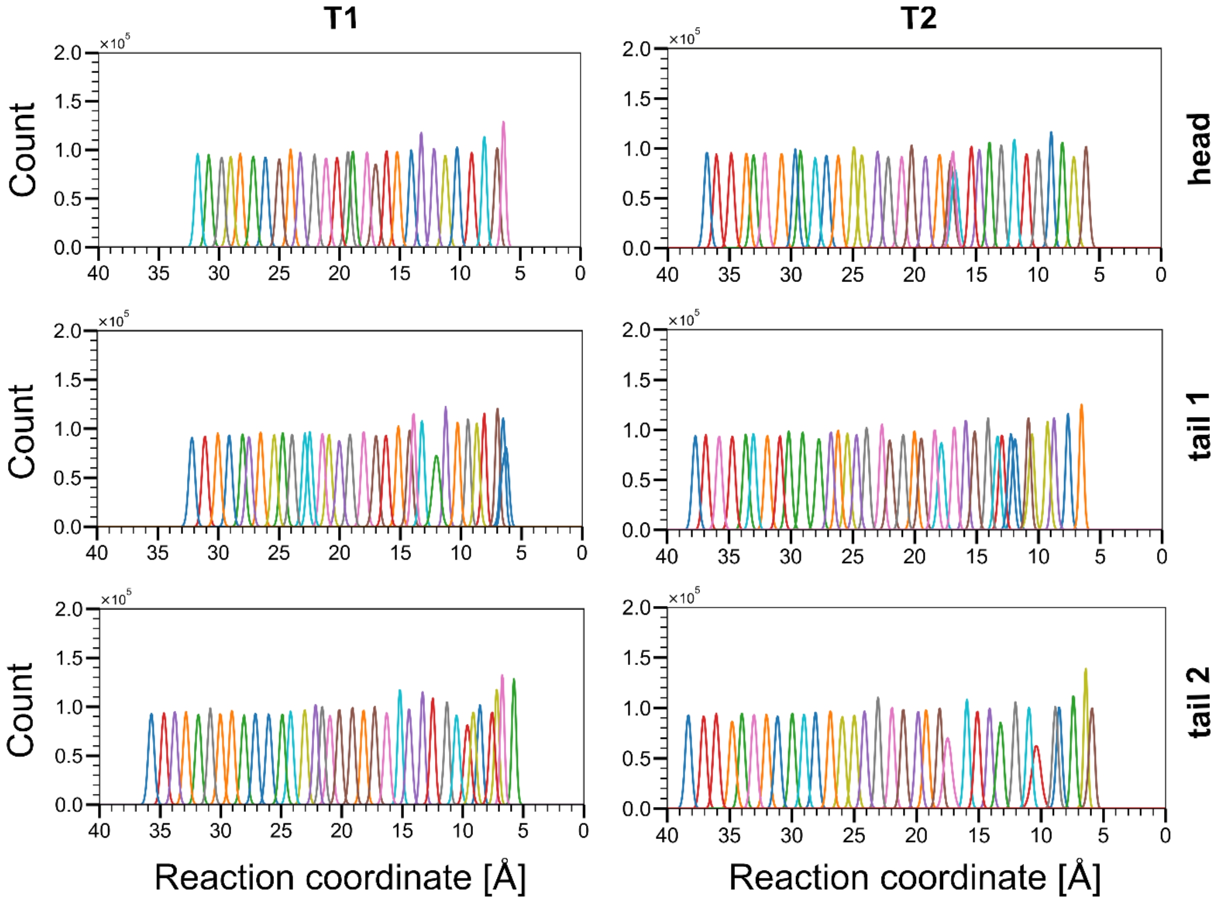


**Figure S9:** Distribution of reaction coordinate values obtained by umbrella sampling for DLPG access via T1 (left) and T2 (right). A force constant of 5 kcal $mol^{-1}$ $Å^{-2}$ was used to restrain the positions of DLPG to the reference point of an umbrella window, which resulted in distributions with a median overlap of at least 4.84% and 3.47% for T1 and T2, respectively.

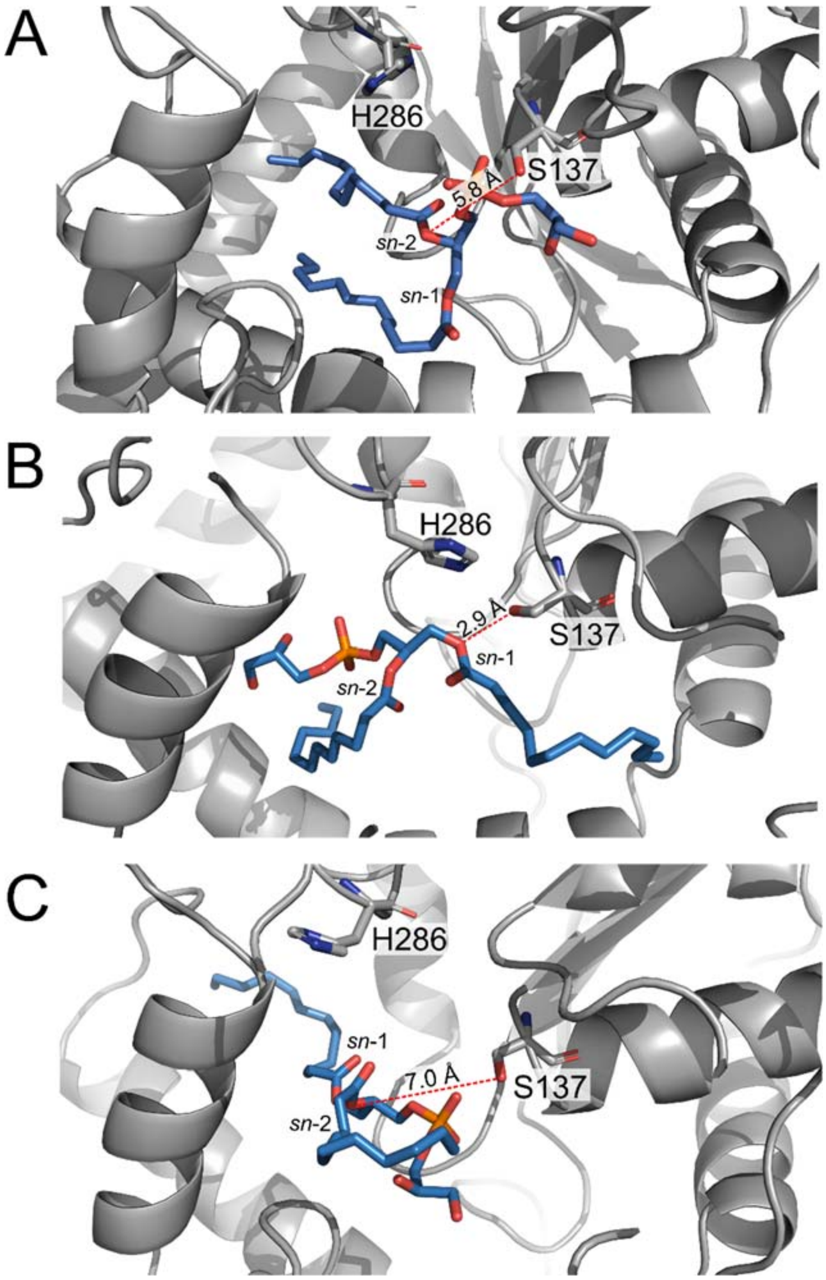


**Figure S10: DLPG access to the catalytic site of t-PlaF$_A$.** DLPG accessing t-PlaF$_A$ through T2 via (A) head first, (B) tail 1 first, and (C) tail 2 first. Snapshots were retrieved after 300 ns of umbrella sampling at the reference point where the substrate's cleavage site is closest to the active site of t-PlaF$_A$. For tail 1, the *sn*-1 site of DLPG comes closest to the catalytic residues (shown in sticks), compared to other access modes.

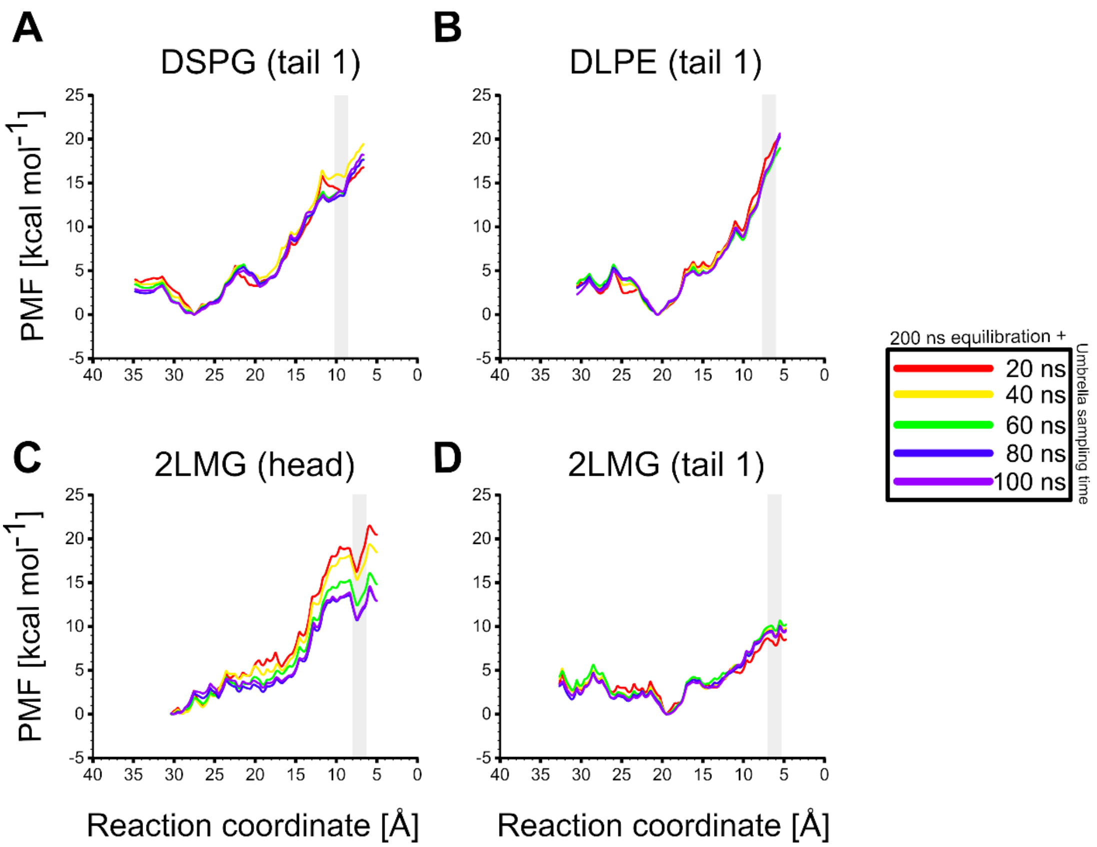


**Figure S11: Convergence of PMFs for other substrate access via T2.** PMFs were computed every 20 ns for the range of 220-300 ns (see legend) of umbrella sampling simulations per window for DSPG-tail 1 access (A), DLPE-tail 1 access (B), 2LMG-head access (C), and 2LMG-tail access (D). The first 200 ns of the sampling simulations were considered for equilibration and removed for every system. The location of the active site is indicated by a grey box. Overall, 300 ns of umbrella sampling per window are sufficient to achieve converged PMFs.

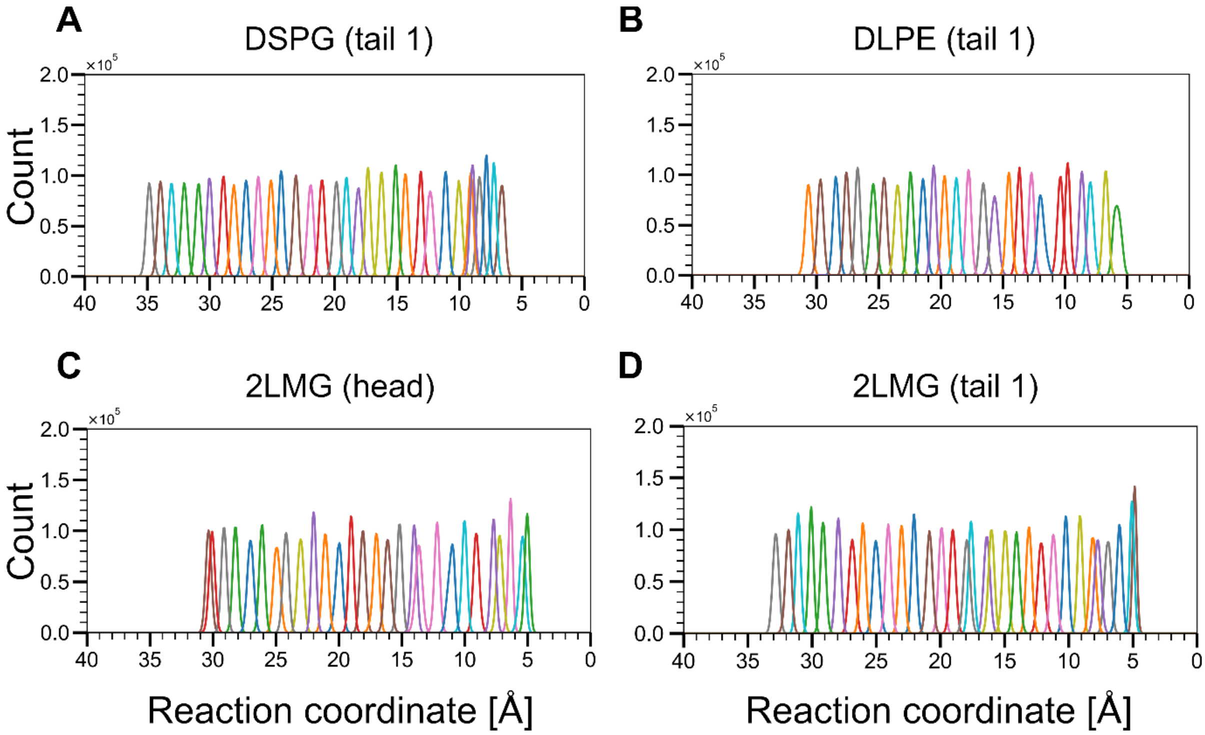


**Figure S12:** Distribution of reaction coordinate values obtained by umbrella sampling for loading of other substrates across T2. A force constant of 5 kcal mol$^{-1}$ Å$^{-2}$ was used to restrain the positions of substrate for (A) DSPG-tail 1 access, (B) DLPE-tail 1 access, (C) 2LMG-head access, and (D) 2LMG-tail 1 access to the reference point of an umbrella window, which resulted in distributions with a median overlap of at least 3.24%.

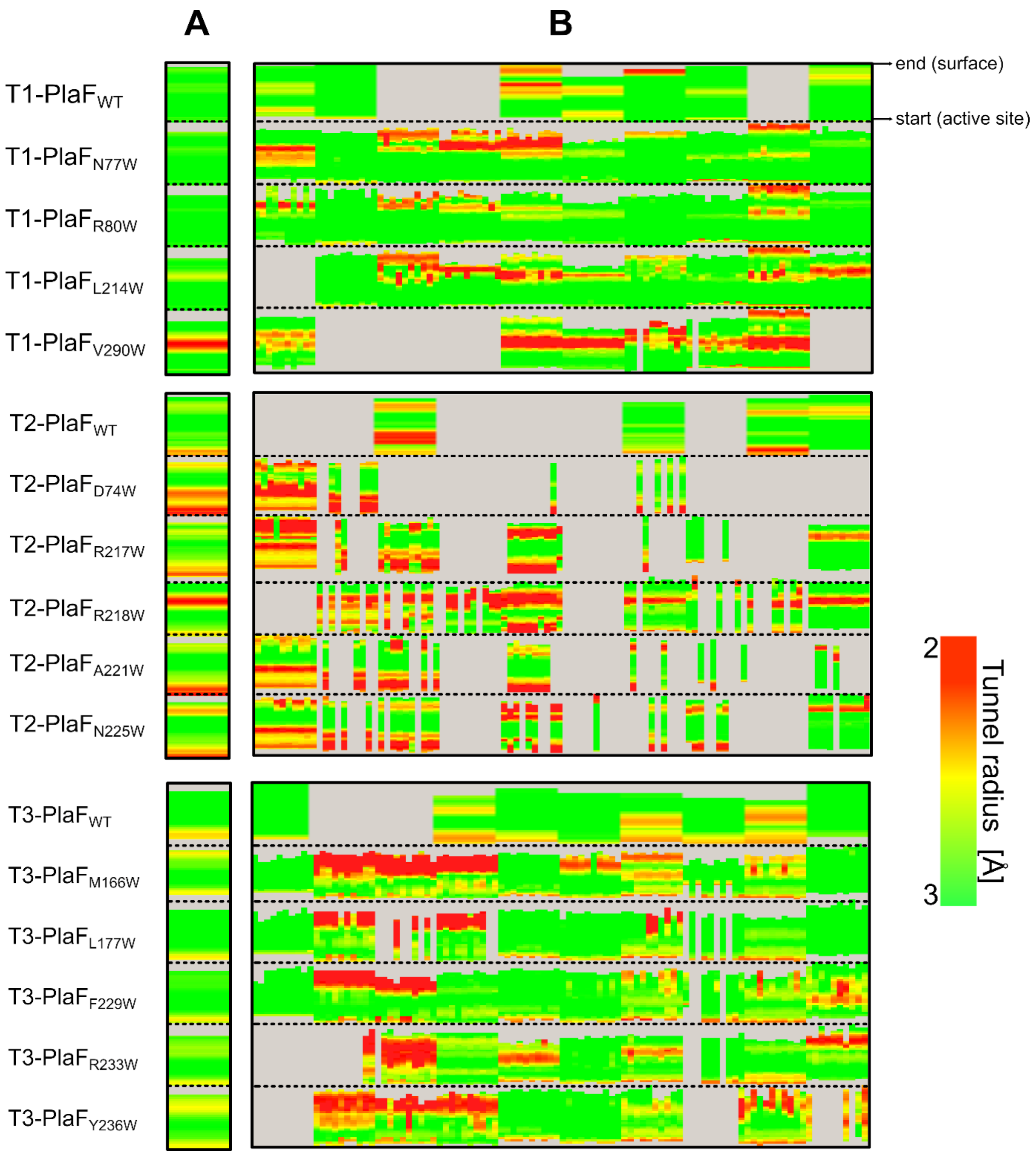


**Figure S13: Heat map visualizing the time evolution of the tunnel profile for the proposed PlaF variants with tryptophan substitutions in T1-T3**. The first row represents the tunnel profile for $PlaF_{WT}$, evaluated from 10 snapshots, obtained at every 200 ns of 2 µs long unbiased MD simulations. Corresponding PlaF variants (second row onwards) were modeled 10 times for each of the 10 snapshots, resulting in 100 snapshots. (A) Average profile for each variant. (B) Time evolution of each variant, with each column corresponding to one snapshot. For $PlaF_{WT}$, continuous snapshots correspond to the increasing time scale of 2 µs in steps of 200 ns. For other variants, every 10 snapshots represent a block of 10 individual profiles of models, obtained from a single snapshot of $PlaF_{WT}$. Each block appears in the increasing time scale of 2 µs. A grey column indicates that the given tunnel was not identified in that particular snapshot. The color scale depicts the tunnel radius.

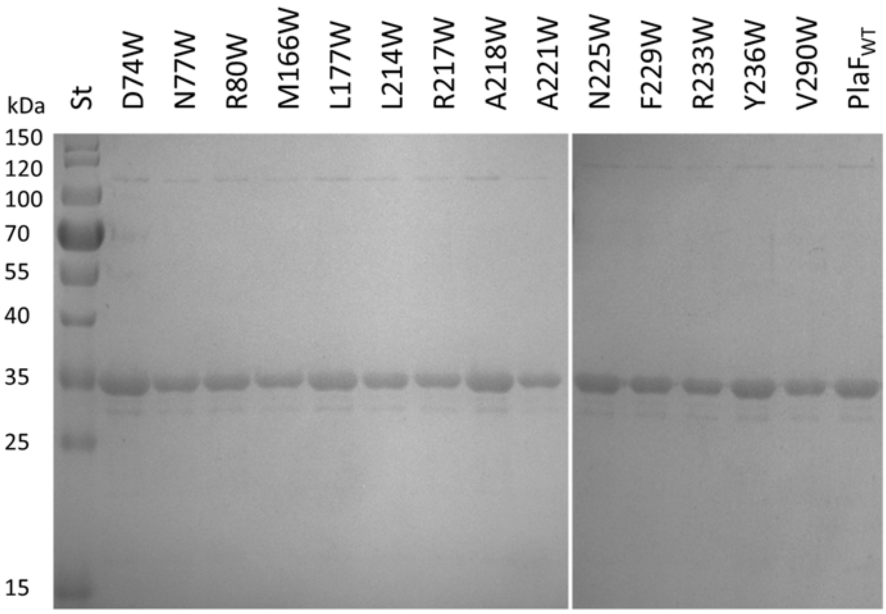


**Figure S14: SDS-PAGE analysis of purified PlaF and variants.** In the Coomassie Brilliant Blue G250-stained gel (14% v/v), PlaF is migrating as ~35 kDa protein. Molecular weights of standard proteins (St) are indicated on the left.

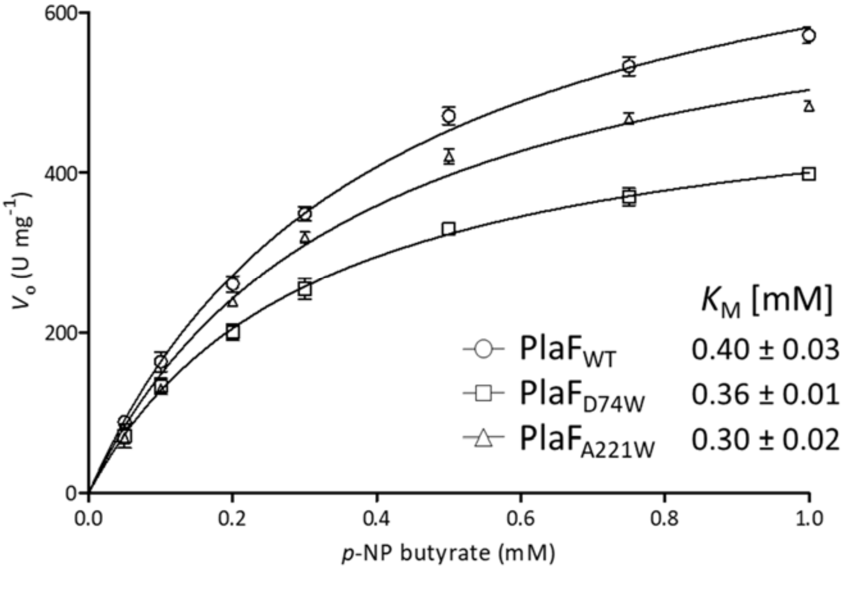


**Figure S15: Enzyme kinetics of the PlaF$_{WT}$ and the substrate-binding tunnel variants.** The activity of PlaF$_{WT}$ and the variants PlaF$_{D74W}$ and PlaF$_{A221W}$ (8 nM) was measured using *p*-NPB, and kinetic parameters were determined by non-linear regression analysis of data fitted to the Michaelis-Menten equation with PrismLab. The results represent the mean ± standard deviation of three independent experiments, $n$ = 9 for PlaF$_{WT}$ and PlaF$_{A221W}$, $n$ = 6 for PlaF$_{D74W}$.

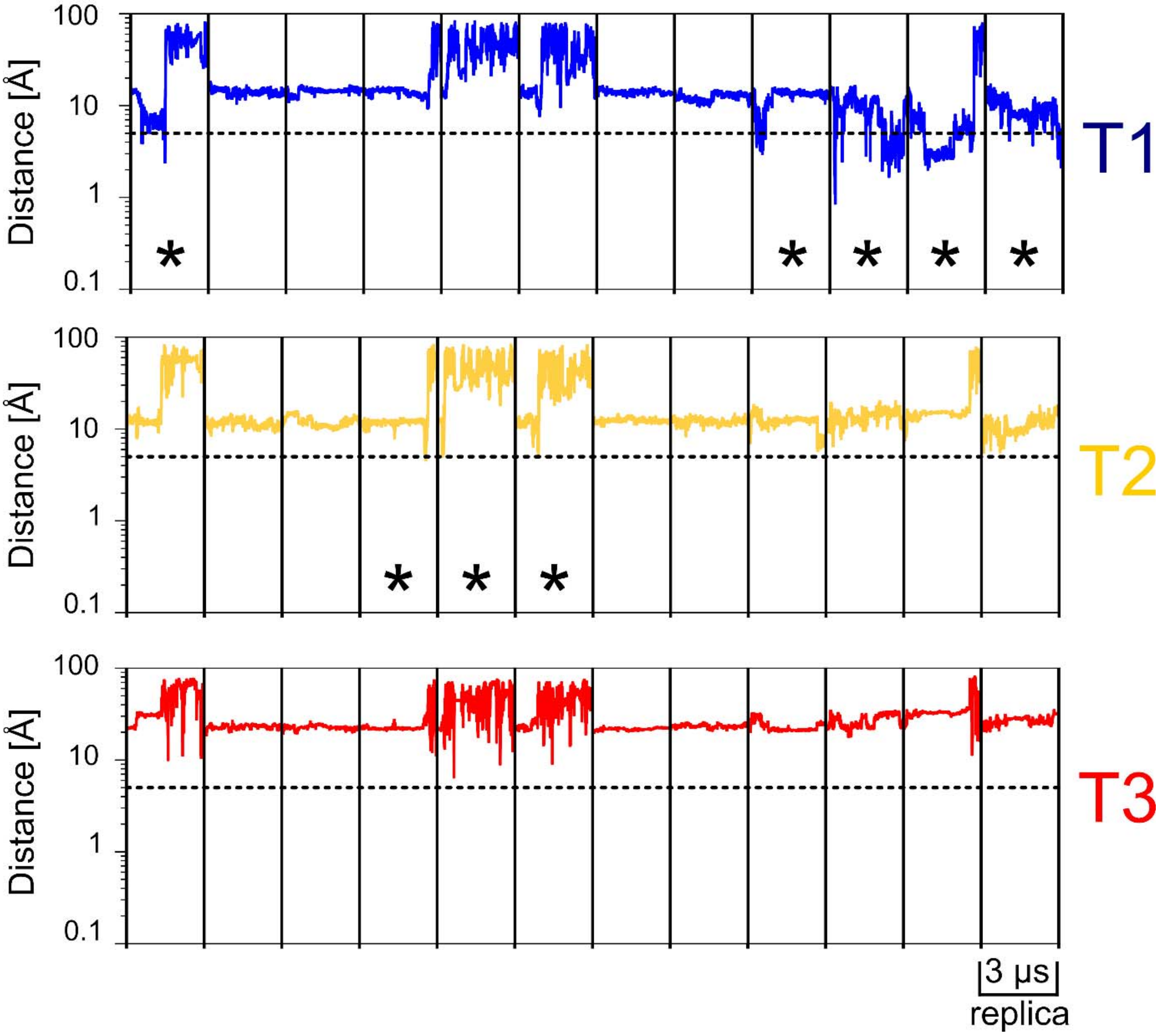


**Figure S16: Unbiased MD simulations of hydrolysis products inside of t-PlaF$_A$.** The distance (in $\log_{10}$ scale) of PGR to the entrance of each of T1-T3 during 12 replicas of 3 µs unbiased MD simulations is plotted, considering the phosphorous atom of PGR. The dashed black line depicts the chosen cutoff of 5 Å, with replicas that reach this cutoff marked with an asterisk. PGR reaches a distance ≤ 5 Å to the entrance of T1 in 5 replicas, including 2 replicas where PGR ultimately leaves T1 to enter into the solvent. In 3 replicas, PGR comes close to the T2 entrance and exits into the solvent.

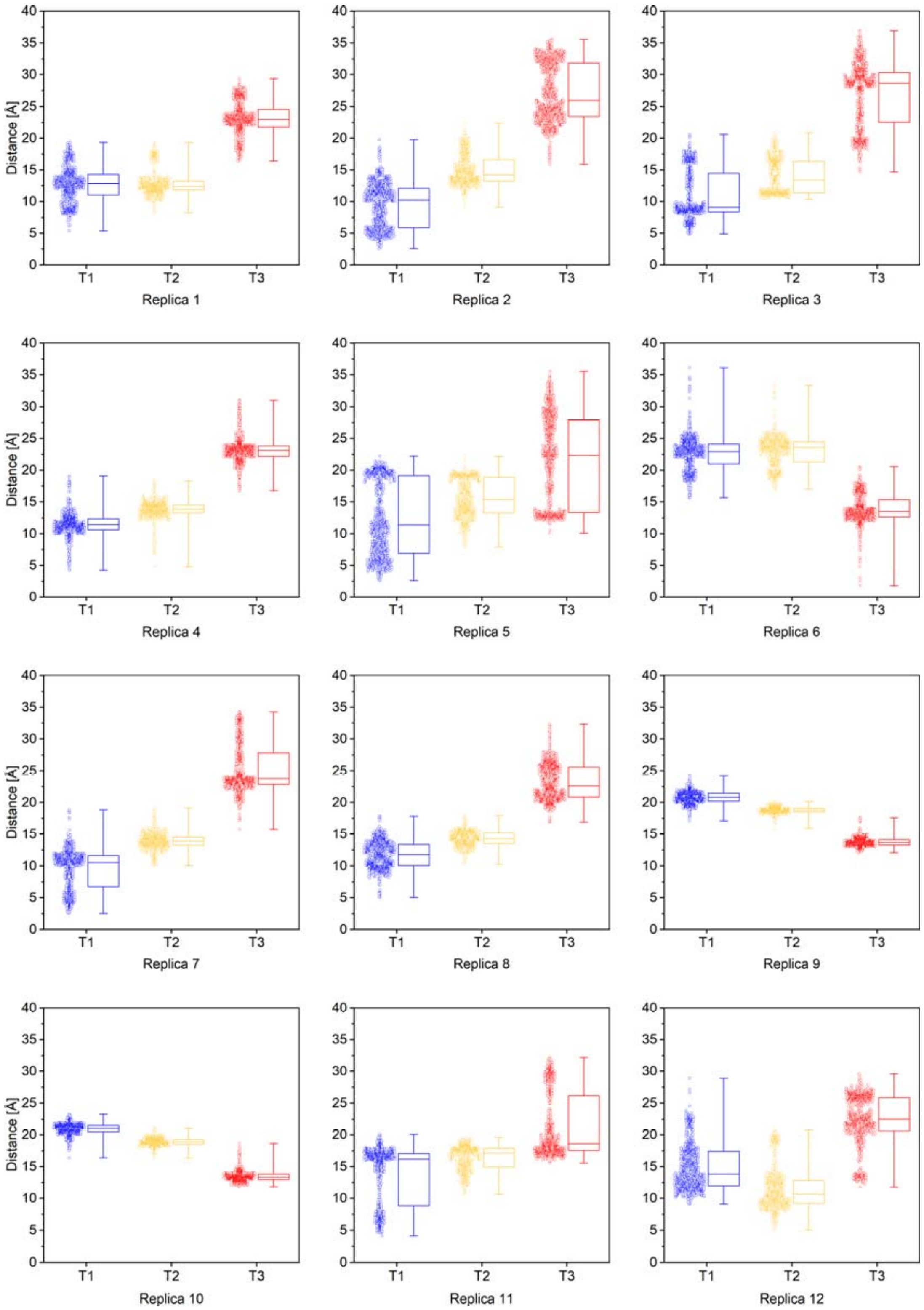


**Figure S17: Unbiased MD simulations of hydrolysis products within t-PlaF$_A$.** For each replica, three box plots represent the distance of MYR to the entrance of tunnels, T1 (blue), T2 (yellow), and T3 (red), during 3 µs long simulations. The box represents the interquartile range between the first and third quartiles, the line inside the box the median, and the whiskers the lowest and highest distance values. The corresponding distribution of data is plotted on the left of each box. Particularly in replica 2, 3, 4, 5, 7, 8, and 11, MYR comes within ~5 Å distance to the entrance of T1, compared to replica 6 where it reaches T3. MYR does not reach T2 at the selected cutoff of 5 Å. These observations indicate that the tendency of MYR reaching the entrance of T1 is significantly higher than for the other two tunnels (Table S6) and likely the egress route for fatty acid products from t-PlaF$_A$.

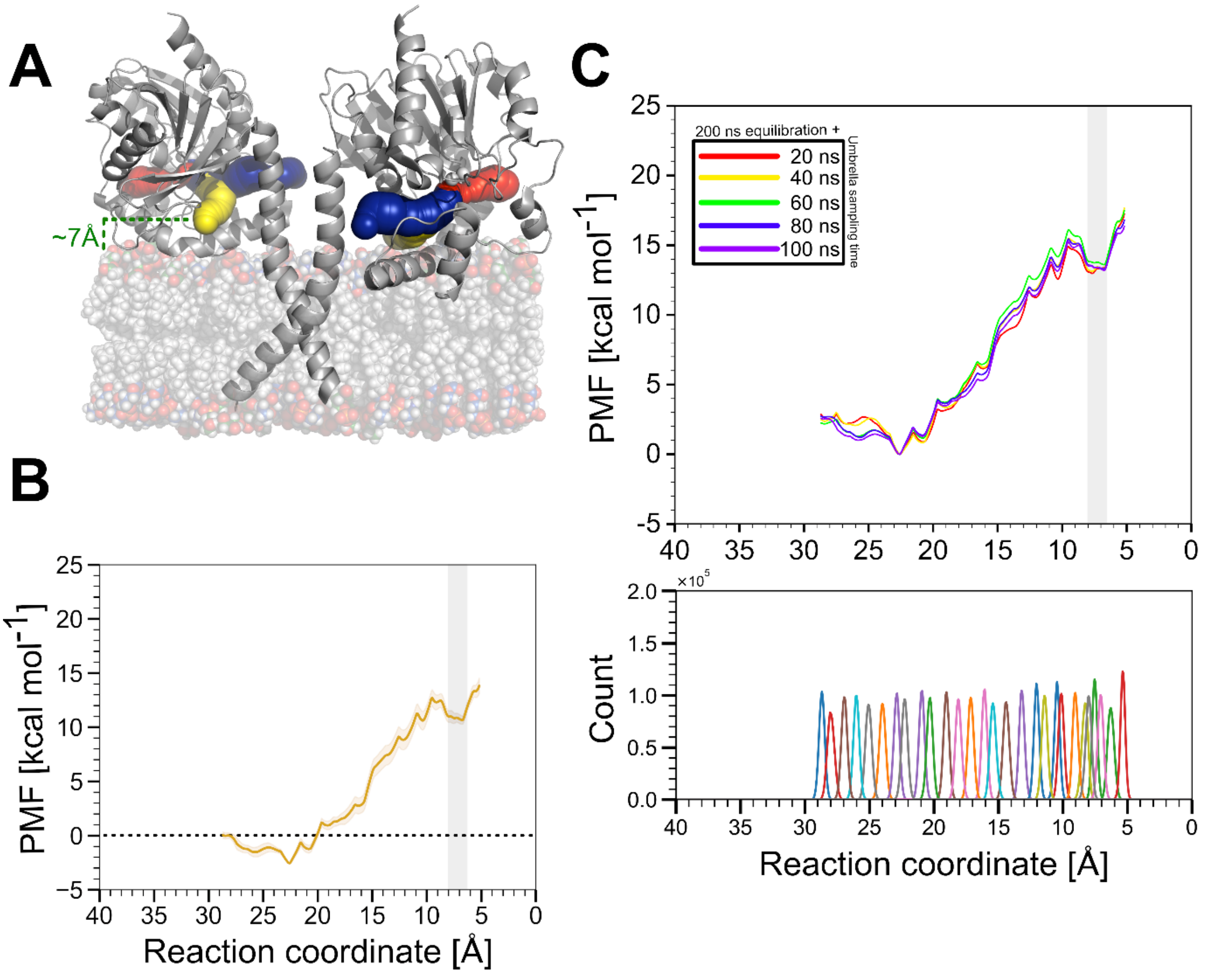


**Figure S18: Substrate access in dimeric PlaF.** A) Among the three tunnels, T2 (yellow spheres) is closest to and its entrance situated at ~7 Å from the membrane interface. The entrances of T1 (blue sphere) and T3 (red sphere) are much farther away (≥ ~12 Å) from the membrane interface, making substrate access into them energetically unfavorable in di-PlaF. B) The PMF of DLPG access via tail 1 across T2 shows a barrier height of 13 kcal $mol^{-1}$ on approaching the catalytic site (grey box). C) Convergence plot (top) indicates sufficient sampling time; the profile converges at 300 ns yielding a maximum difference of ~0.5 kcal $mol^{-1}$ compared to the PMF computed at 280 ns. The histograms (bottom) indicate sufficient overlap among the umbrella windows using a force constant of 5 kcal $mol^{-1}$ $Å^{-2}$; the median overlap is 4.2%.

### 4. Supplementary movies

**Movie S1:** The acyl chain termini of lipids can reach the membrane interface in the presence of t-PlaF$_A$ during 40 ns long unbiased MD simulation. Grey cartoon represents t-PlaF$_A$, which is embedded in the membrane (grey sticks). The termini of acyl chains of lipids in the membrane are shown as red spheres. At the beginning of the simulation, an acyl terminus of the selected lipid (blue spheres) stretches towards the center of the membrane and then moves towards the membrane interface at around 15 ns and stays there for ~10 ns, until it comes back to the bilayer center (see Figure S3A, black curve in II for Memb + t-PlaF$_A$).

**Movie S2:** The acyl chain termini of lipids can reach the membrane interface in the absence of t-PlaF$_A$ during 40 ns long unbiased MD simulation. The Membrane is shown as grey sticks. The termini of acyl chains of lipids in the membrane are shown as red spheres. At the beginning of the simulation, the acyl termini of the selected lipid (blue spheres) remains at the center of the membrane and then after 19 ns one of the tail starts moving towards the membrane interface (see Figure S3A, black curve in II for Memb - t-PlaF$_A$).

**Movie S3:** Extraction of a DLPG substrate molecule via the head from the membrane into T1 of t-PlaF$_A$ using sMD simulations. Grey cartoon represents t-PlaF$_A$, which is embedded in the membrane composed of 70% DLPE (grey spheres) and 30% DLPG (black spheres). DLPG (blue spheres) is pulled via the head. The *sn*-1 and *sn*-2 sites of the pulled DLPG are shown as orange and magenta spheres, respectively. During the course of the simulations, the *sn*-2 site of DLPG reaches close to the catalytic S137 (red sticks).

**Movie S4:** Extraction of a DLPG substrate molecule via the tail 1 from the membrane into T1 of t-PlaF$_A$ using sMD simulations. Grey cartoon represents t-PlaF$_A$, which is embedded in the membrane composed of 70% DLPE (grey spheres) and 30% DLPG (black spheres). DLPG (blue spheres) is pulled via the tail 1. The *sn*-1 and *sn*-2 sites of the pulled DLPG are shown as orange and magenta spheres, respectively. During the course of the simulations, the *sn*-1 site of DLPG reaches close to the catalytic S137 (red sticks).

**Movie S5:** Extraction of a DLPG substrate molecule via the tail 2 from the membrane into T1 of t-PlaF$_A$ using sMD simulations. Grey cartoon represents t-PlaF$_A$, which is embedded in the membrane composed of 70% DLPE (grey spheres) and 30% DLPG (black spheres). DLPG (blue spheres) is pulled via the tail 2. The *sn*-1 and *sn*-2 sites of the pulled DLPG are shown as orange and magenta spheres, respectively. During the course of the simulations, the *sn*-2 site of DLPG reaches close to the catalytic S137 (red sticks).

**Movie S6:** Extraction of a DLPG substrate molecule via the head from the membrane into T2 of t-PlaF$_A$ using sMD simulations. Grey cartoon represents t-PlaF$_A$, which is embedded in the membrane composed of 70% DLPE (grey spheres) and 30% DLPG (black spheres). DLPG (blue spheres) is pulled via the head. The *sn*-1 and *sn*-2 sites of the pulled DLPG are shown as orange and magenta spheres, respectively. During the course of the simulations, the *sn*-2 site of DLPG reaches close to the catalytic S137 (red sticks).

**Movie S7:** Extraction of a DLPG substrate molecule via the tail 1 from the membrane into T2 of t-PlaF$_A$ using sMD simulations. Grey cartoon represents t-PlaF$_A$, which is embedded in the membrane composed of 70% DLPE (grey spheres) and 30% DLPG (black spheres). DLPG (blue spheres) is pulled via the tail 1. The *sn*-1 and *sn*-2 sites of the pulled DLPG are shown as orange and magenta spheres, respectively. During the course of the simulations, the *sn*-1 site of DLPG reaches close to the catalytic S137 (red sticks).

**Movie S8:** Extraction of a DLPG substrate molecule via the tail 2 from the membrane into T2 of t-PlaF$_A$ using sMD simulations. Grey cartoon represents t-PlaF$_A$, which is embedded in the membrane composed of 70% DLPE (grey spheres) and 30% DLPG (black spheres). DLPG (blue spheres) is pulled via the tail 2. The *sn*-1 and *sn*-2 sites of the pulled DLPG are shown as orange and magenta spheres, respectively. During the course of the simulations, the *sn*-2 site of DLPG reaches close to the catalytic S137 (red sticks).
